# Air and environmental sampling expand metagenomic viral surveillance in Cambodian pig slaughterhouses

**DOI:** 10.64898/2026.11.16.765117

**Authors:** Peter Cronin, Vireak Heang, Jurre Y. Siegers, Songha Tok, Sarath Sin, Ben Sievers, Victor Omondi, Sithun Nuon, Kimtuo Chhel, Janin Nouhin, Vutha Chim, Bunnary Seng, Makara Hak, Sorn San, Sothyra Tum, Filip F. Claes, Cadhla Firth, Yvonne C.F. Su, Gavin J.D. Smith, Erik A. Karlsson

## Abstract

Direct animal sampling describes selected hosts but can miss the wider environment through which viral material is shed, mixed and transported. Across nine visits to two Cambodian pig slaughterhouses, we applied capture-enriched metagenomic sequencing to air samples (n = 25), faeces (n = 156), wastewater (n = 15), carcass wash water (n = 18), holding-pen swabs (n = 124), slaughter-box swabs (n = 73) and ropes (n = 16), comparing viral detections with those concurrently collected from pig snout swabs (n = 193). Combined environmental sampling recovered 96.6% of the viral taxa detected in snout swabs overall, exceeding 90.9% at every visit, while also detecting additional swine-associated taxa as well as viral RNA associated with avian and human hosts. Air alone provided the widest coverage from a single environmental sample type, recovering more than 77% and retaining many common swine-associated viruses and contributing the largest number of detections not recovered from any other environmental sample. Other samples captured different shedding routes and stages of slaughter. Slaughter-box swabs recovered the fewest taxa, showing that visible exposure to animal material did not guarantee surveillance breadth. Air expanded the reach of non-contact surveillance, but the slaughterhouse became most visible when several environmental sample types were interpreted together.

## Introduction

Global pig production continues to expand, and growing international trade and animal movement increase the opportunities for viruses to spread from and between pigs. A slaughterhouse is a critical point within these production chains. It aggregates animals from multiple farms, brings workers and animals into close contact, and redistributes biological material through air, water, surfaces and waste. This matters because pigs maintain and exchange viruses with both animal and human populations, and several viruses of One Health importance illustrate this. Porcine deltacoronavirus (PDCoV, also known as coronavirus HKU15), an enteric coronavirus of pigs, has infected children in Haiti through at least two independent zoonotic events [1]. Pigs have also acted as bridging hosts during Nipah virus outbreaks [2, 3] and as amplifying hosts for Japanese encephalitis virus [4–7], and they remain important hosts for influenza A viruses [8], including the swine lineages from which the 2009 A(H1N1) pandemic virus emerged [9, 10]. Pigs are also the principal animal reservoir of hepatitis E virus genotypes 3 and 4, which reach people through undercooked pork and through occupational contact in farming and slaughter settings [11, 12]. Surveillance is therefore most valuable where animal production, movement, slaughter and occupational exposure converge [13].

The same interface matters for animal health and food production. African swine fever virus, porcine epidemic diarrhoea virus, porcine reproductive and respiratory syndrome virus and classical swine fever virus have caused major disruption across swine-producing regions [14–24]. Slaughterhouses offer a practical surveillance point because they consolidate animals from multiple farms and provinces without requiring investigators to visit each source herd. Repeated sampling at slaughter can therefore describe part of a supply network rather than a single farm [25, 26]. The difficulty is deciding what, exactly, should be sampled.

Conventional swine-virus surveillance relies largely on direct animal sampling, followed by targeted molecular assays, antigen detection and, less commonly, sequencing. These approaches are essential when the question concerns an individual animal, but they require animal handling, trained personnel, cold-chain logistics and sufficient sampling density to capture heterogeneous circulation. They are also anatomically selective. A snout swab can miss viruses shed through non-respiratory routes and material deposited elsewhere in the facility. Precision at the level of one animal can therefore leave the wider surveillance interface poorly described.

Environmental sampling paired with capture-enriched metagenomic sequencing offers another view. Rather than choosing an animal in advance, it samples biological material accumulated from multiple hosts, routes and activities. Detection remains limited to viruses represented in the capture panel and sufficiently similar in sequence to its probes, while viral nucleic acid recovered from an environmental sample does not identify its host of origin or show that the virus was infectious [27]. Air is especially attractive because it can integrate material released by multiple animals and activities without direct handling. The unresolved question is whether the viruses detected in air represent the viruses circulating through the facility, a selective fraction of them, or mostly environmental background. Environmental metagenomics has detected circulating viruses in high-risk animal settings, including highly pathogenic avian influenza A(H5N1) virus in Cambodian live-poultry markets [27], but comparisons among air, other environmental samples and pig samples remain limited [28].

Here, viral detections from air, faeces, wastewater, carcass wash water, holding-pen swabs, slaughter-box swabs and ropes were compared with detections from pig snout swabs collected during the same visits to two Cambodian slaughterhouses. The study asked three practical questions: how much of the snout-swab virus composition could environmental sampling recover, what did air add, and which viruses would have been missed had only one environmental sample type been collected?

## Materials and methods

### Sample collection

This study was conducted between November 2022 and March 2023 at two swine slaughterhouses in Cambodia: one on the outskirts of Doun Kaev, Takeo Province, and one in Phnom Penh, the country’s largest urban centre (Supplementary Fig. 1). Nine visits were completed, comprising six in Phnom Penh and three in Takeo (Supplementary Table 1). Poultry slaughter was observed once at the Phnom Penh facility, within the same open area and sharing the same air volume and drainage as the pigs. Because each visit was brief, the frequency of poultry slaughter at either facility could not be established, and poultry were not directly sampled.

At each visit, environmental samples and pig snout swabs were collected at the same slaughterhouse on the same day. All occupied holding pens accessible during the visit were sampled (Supplementary Table 1); no fixed set of pens was followed over time. Snout swabs, oral fluids and faeces were collected from the same groups of pigs. Cotton ropes were suspended in each pen for 180 min to collect oral and nasal secretions deposited through chewing, rubbing and direct contact, as previously described [29]. Fresh faeces were collected from the pen floor, and pen surfaces were swabbed while occupied. All swab-based samples were collected using polyester swabs placed in in-house viral transport medium (VTM), as previously described [27]. Individual snout swabs were placed in 1.5 mL of VTM and stored at Institut Pasteur du Cambodge until processing; equal volumes (25 µL) from ten swabs collected in a pen were pooled immediately before RNA extraction.

Adjacent holding pens were separated by metal railings that permitted direct contact between animals. The holding area was therefore treated as one epidemiological unit rather than a set of independent pens. Pen identifiers were not recorded for individual snout-swab pools, faecal samples or pen-surface swabs. Although these samples were collected from the same holding area on the same day, individual samples could not be attributed to a specific pen. Pig and environmental samples were therefore matched by slaughterhouse and sampling date. Wastewater, carcass wash water and slaughter-box swabs were collected at the facility level and were not linked to individual pens. Each slaughterhouse contained one slaughter box. At each visit, 50 mL of carcass wash water and 50 mL of wastewater were collected.

Air was sampled at three positions per slaughterhouse per visit using AerosolSense samplers (Thermo Fisher Scientific), as previously described [27]. Samplers operated at 200 L/min through an omnidirectional inlet for 180 min. Two samplers were positioned inside or immediately beside the sampled holding pens, and one was positioned approximately 30 m from the holding area. Material from each sampler cartridge was eluted in 2 mL of VTM, and each cartridge was extracted and sequenced separately. All samples were transported on ice to Institut Pasteur du Cambodge in Phnom Penh, aliquoted and stored at -80 °C until processing.

### RNA extraction

Snout swabs were pooled in groups of up to ten animals, yielding 20 pools from 193 animals. Environmental samples of the same type collected at the same visit were pooled in the same way (up to ten samples per pool), with the exception of air samples, which were processed individually. In total, 620 individual samples were combined into 103 sequencing libraries: 20 from snout swabs and 83 from environmental samples (427 individual samples) (Supplementary File 1). RNA was extracted using the Direct-zol RNA Miniprep Kit (Zymo Research, California, USA) according to the manufacturer’s protocol. A 200-µL aliquot was mixed with TRI Reagent or TRIzol at a 1:3 ratio. An equal volume of absolute ethanol was added, and the mixture was loaded onto a Zymo-Spin IICR column. Columns were washed twice using Direct-zol RNA PreWash followed by RNA Wash, and RNA was eluted in 50 µL of nuclease-free water. Extracts were taken forward to library preparation without DNase treatment, so both DNA and RNA viral genomes were retained. RNA templates additionally required reverse transcription and second-strand synthesis, as described below, whereas DNA templates entered library preparation directly.

### Library preparation and sequencing

Sequencing libraries were generated using the Twist Library Preparation Enzymatic Fragmentation Kit 2.0 and Twist Comprehensive Viral Research Panel (Twist Bioscience, San Francisco, California, USA) according to the manufacturer’s instructions. RNA was reverse transcribed using the ProtoScript II First Strand cDNA Synthesis Kit with Random Primer 6 (S1230S; New England Biolabs, Ipswich, Massachusetts, USA), followed by second-strand synthesis using the NEBNext Ultra II Non-Directional RNA Second Strand Synthesis Kit (E6111S). Libraries compatible with Illumina TruSeq chemistry were prepared using the Twist EF Kit 2.0, with fragmentation at 37 °C for 5 min and 10 cycles of pre-capture PCR. Target enrichment used the Twist Comprehensive Viral Research Panel, followed by 22 cycles of post-capture PCR. Final libraries were pooled in equimolar ratios and sequenced on an Illumina MiSeq using a v3 reagent kit to generate paired-end 75-bp reads.

### Processing of raw viral sequencing data and abundance estimation

Raw sequencing reads were processed using the SeqtoID pipeline, formerly CZ ID, as previously described [30]. Read counts mapping directly to de novo assembled viral contigs were downloaded after processing. Taxa associated with fungi, plants or bacteria, including members of the class Caudoviricetes and order Crassvirales, were excluded. Viral abundance was expressed as reads per million (RPM), calculated by dividing reads assigned to each viral species by the total reads in the corresponding library and multiplying by 10⁶. To reduce potential false-positive assignments, reads mapping to viral contigs were retained only when they comprised more than 0.1% of total reads in at least one sample; values that did not meet this criterion were set to zero.

### Virus diversity and compositional similarity estimation

For beta-diversity analyses, viral read-count data were transformed using vegdist in vegan v2.7-3 [31] to calculate Jaccard or Bray-Curtis dissimilarity. Principal coordinate analysis (PCoA) visualised beta-diversity, and PERMANOVA tested overall compositional differences using adonis2. Pairwise comparisons used pairwise.adonis. Spearman correlations between virus detection frequency and distance from median PCo1 and PCo2 coordinates were calculated using corr.test in psych v2.6.3. To estimate the contribution of individual species to compositional differences between pig and environmental samples, the mean Jaccard distance between samples collected at the same time point was calculated. Each viral taxon was then removed sequentially, Jaccard distance was recalculated, and the change in distance quantified that taxon’s contribution. Mean changes across sampling dates summarised overall contributions. Principal components analysis used prcomp in stats v4.5.2. Alpha-diversity was calculated using estimate_richness in phyloseq v1.54.2 and included observed richness, Chao1, Shannon and Simpson indices.

### Viral sequence similarity and coverage quantification

The top DIAMOND blastx hit was extracted for each viral contig assembled de novo from pig snout-swab samples using the CZ ID pipeline. The resulting 3,701 viral proteins were downloaded from GenBank and used to construct a custom reference database with DIAMOND makedb. Quality-filtered contigs were queried against this database using DIAMOND blastx. Coverage, alignment length, mismatches, percentage identity and e-value were recorded, and alignments with e-values greater than 1 × 10⁻⁵ were excluded. Pig and environmental contigs were compared when both aligned to the same reference gene. Genes without a qualifying alignment in either sample were classified as undetected. Differences in percentage identity were calculated by subtracting the identity of the environmental contig from that of the pig contig.

### Statistics and graphical representation

All statistical analyses were performed in R v4.5.2 [32]. Kruskal-Wallis tests with Dunn’s post hoc comparisons used kwAllPairsDunnTest in PMCMRplus v1.9.12. Comparisons between two groups used Wilcoxon rank-sum tests, and differences in detection frequency between locations used Fisher’s exact tests. P values were adjusted for false discovery using the Benjamini-Hochberg method. Unless otherwise stated, adjusted P values are annotated as * P < 0.05, ** P < 0.01 and *** P < 0.001. Visualisations were generated primarily using ggplot2 v4.0.3. Heatmaps used ComplexHeatmap v2.26.1. For heatmap visualisation, RPM values were rank-scaled to the interval [0,1] using x_i^R = [Rank(x_i) - Minimum(Rank(x))]/[Maximum(Rank(x)) - Minimum(Rank(x))]. This transformation preserved the relative ordering of abundance while reducing the influence of extreme values. Detection overlap was visualised using chordDiagram in circlize v0.4.18 and radarchart in fmsb v0.7.6.

## Data availability

Data generated in this study are available through the European Nucleotide Archive under accession number PRJEB129191.

## Code availability

Analysis scripts, processed data tables and figure-generation code are available through Zenodo (https://doi.org/10.5281/zenodo.23793111) and mirrored on GitHub (https://github.com/yvonnesuLoVE/Pig-Environmental-Metagenomics).

## Results

### Combined environmental sampling recovers most snout-swab detections

The first benchmark was the 88 species-level viral taxa detected in pig snout swabs. Across the study, these represented 378 time point-specific detections, defined as detection of one viral taxon in one sample type on one sampling date (Supplementary Fig. 2; Supplementary Table 2). When all environmental samples were combined, 365 of the 378 snout-swab detections (96.6%) were also recovered environmentally, and recovery exceeded 90.9% at every visit (Fig. 1a; Supplementary Fig. 2a). Only 13 of the 378 snout-swab detections (3.4%) were not recovered environmentally. These included Pestivirus K, porcine mastadenovirus A, porcine stool-associated circular virus 5 and porcine circovirus 2 (Fig. 1a).

**Figure 1.**
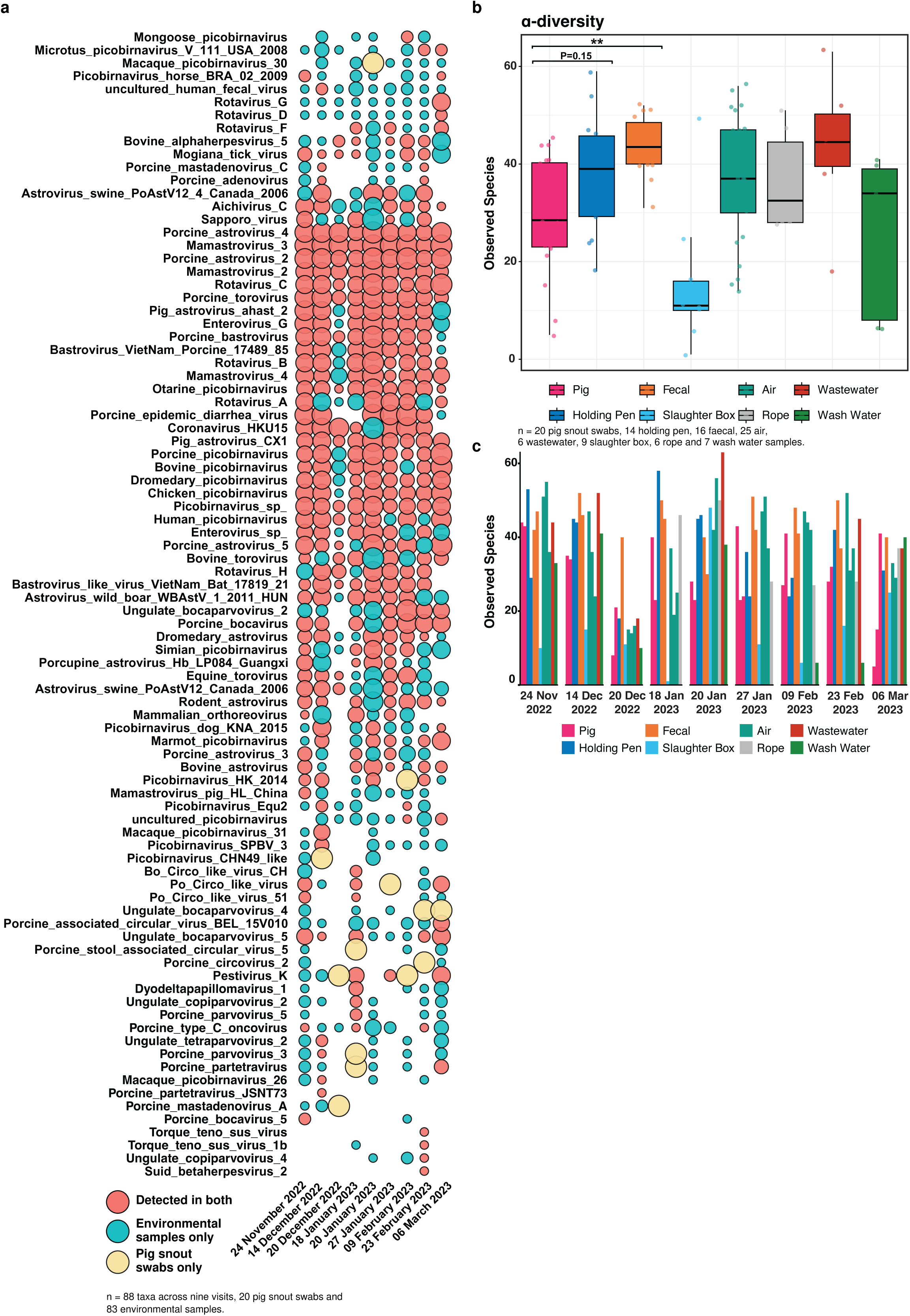
Combined environmental sampling captures most viral taxa detected in pig snout swabs. a, Detection status of 88 viral taxa across nine visits (six to Phnom Penh and three to Takeo). Bubble colour indicates whether a taxon was detected in both environmental samples and pig snout swabs (red), in environmental samples only (teal), or in pig snout swabs only (yellow), and bubble size reflects the number of samples in which the taxon was detected. b, Observed viral richness across sample types (20 pig snout swabs, 14 holding pen, 16 faecal, 25 air, 6 wastewater, 9 slaughter box, 6 rope and 7 wash water samples). Box plots show the median, interquartile range and range, and points show individual samples. Differences between sample types were tested using Kruskal-Wallis tests with Dunn’s post hoc comparisons and Benjamini-Hochberg correction. Adjusted P values are annotated as * P < 0.05, ** P < 0.01 and *** P < 0.001. c, Observed viral richness by sample type and sampling date.

The environmental profile was not simply a copy of the snout-swab profile (Fig. 1a; Supplementary Fig. 3b). For example, on 24 November 2022, snout swabs contained 49 viral taxa and environmental samples contained 79, while on 20 January 2023 environmental sampling detected 72 taxa, twice the number found in snout swabs. Environmental samples contained more taxa than snout swabs at all nine visits (Supplementary Fig. 3b). Air contributed 44 detections that were absent from snout swabs and from every other environmental sample type on the same visit, the largest contribution from one sample type, followed by faeces with 38. Each remaining sample type contributed between only five and eight (Supplementary Fig. 4a-b).

Viral richness varied sharply among sample types. Faecal samples contained significantly more observed viral species than snout swabs (P = 0.008), whereas all other environmental sample types did not differ significantly from snout swabs in observed richness, Chao1 or Shannon index (Fig. 1b,c; Supplementary Fig. 5; Supplementary Tables 3-5). Slaughter-box swabs contained the fewest taxa and were significantly less diverse than most other environmental sample types (Fig. 1b,c; Supplementary Table 3).

A large overlap also did not imply identical virus composition. Jaccard PCoA separated sample types after accounting for location and sampling date (PERMANOVA, FDR-adjusted P < 0.001, R² = 0.15; Fig. 2a). Holding-pen swabs, slaughter-box swabs and faeces differed significantly from snout swabs, whereas ropes, wastewater and carcass wash water did not (Supplementary Fig. 6a; Supplementary Tables 6 and 7). Air also differed from snout swabs, with much of the separation occurring along PCo2 (Supplementary Fig. 6b; Supplementary Tables 6 and 8).

**Figure 2.**
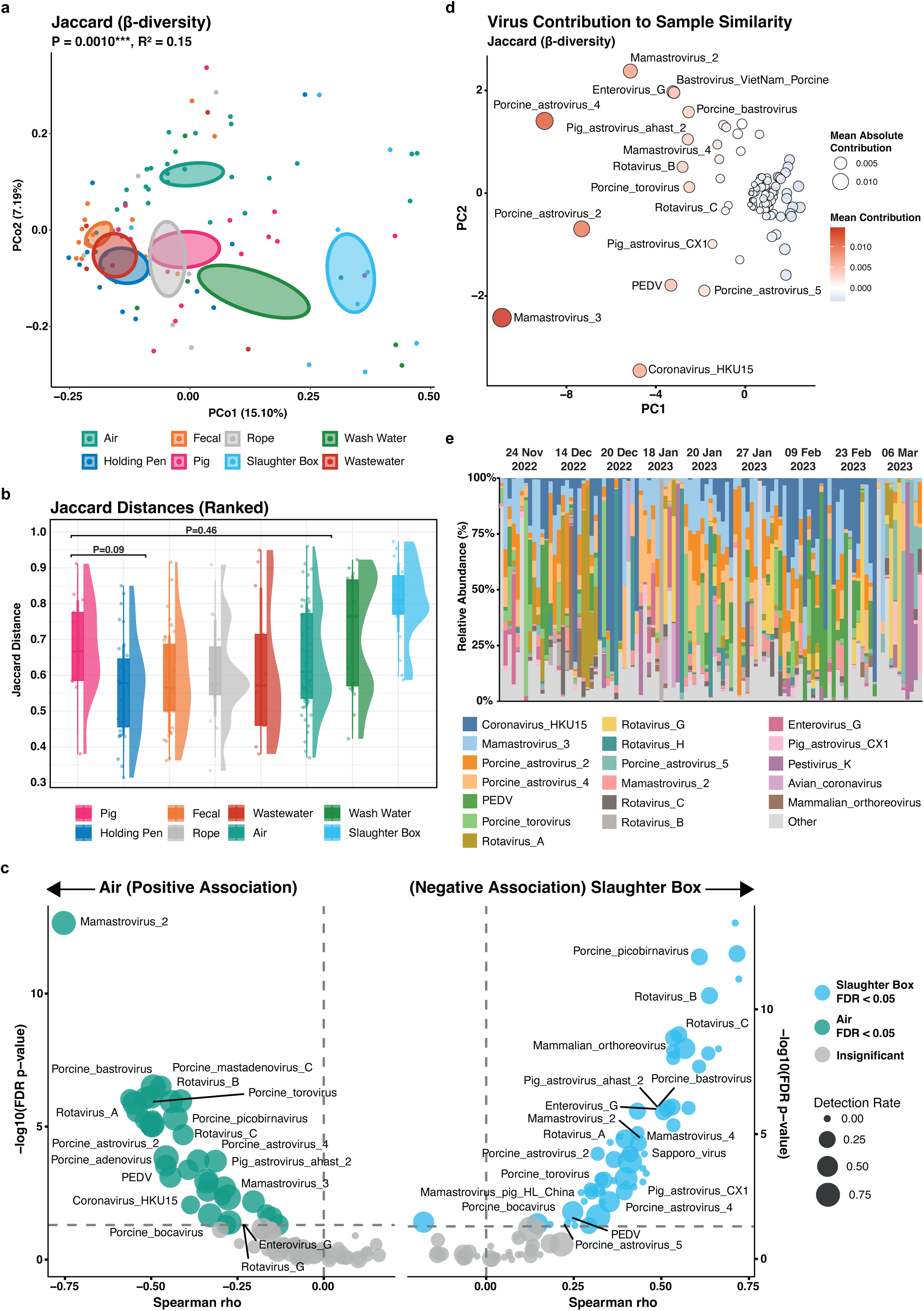
Environmental sample types capture distinct virus composition. a, Principal coordinate analysis of Jaccard dissimilarity among sample types (n = 103 samples). PERMANOVA tested differences in virus composition among sample types. b, Jaccard distance between pig snout swabs and environmental samples collected at corresponding visits. c, Spearman correlations between virus detection frequency and distance from the median PCoA coordinates for air and slaughter box samples. d, Principal components analysis of the contribution of individual taxa to similarity between sample types, estimated through sequential removal and recalculation of Jaccard distance. e, Relative abundance of the most abundant viral taxa across sample types and visits. P values were adjusted for false discovery using the Benjamini-Hochberg method.

### Different environmental samples expose different parts of the slaughterhouse

Jaccard distance from snout swabs collected at the same place and time provided one measure of what each sample type captured (Fig. 2b). Holding-pen swabs were most similar to snout swabs, followed by faeces, ropes, wastewater and air. Carcass wash water and slaughter-box swabs were more distant, consistent with material accumulating differently across the slaughter process (Supplementary Table 9). No individual sample type reproduced the combined environmental profile. Each opened a different window onto hosts, shedding routes, activities and locations within the facility.

### Air provides the widest coverage from a single environmental sample type

Air was compositionally distinct from snout swabs but retained many common swine-associated viruses and contributed more sample-specific detections than any other environmental sample type. Detection of 37 swine-associated taxa correlated negatively with distance from the median air centroid, including coronavirus HKU15, porcine epidemic diarrhoea virus and members of the Astroviridae, Adenoviridae and Tobaniviridae (Fig. 2c; Supplementary Table 10). Many also correlated negatively with distance from the snout-swab centroid (Supplementary Fig. 7; Supplementary Table 11). Air did not simply reproduce the virus composition of selected pigs. Within the present sampling design, it integrated material generated by multiple animals, shedding routes and activities over a three-hour period. Carcass wash water showed the same direction of association for fewer taxa, with 23 of 26 significant taxa correlating negatively with distance from the carcass wash water centroid (Supplementary Table 12).

### The slaughter box is a poor generalist but a useful counterexample

The slaughter box looked like an obvious surveillance surface but recovered few taxa. Detection of 55 of the 56 taxa reaching significance correlated positively with distance from the slaughter-box centroid, consistent with limited recovery of the wider swine-associated virus composition (Fig. 2c; Supplementary Table 13). Yet poor general coverage did not mean zero value. Slaughter-box swabs repeatedly detected coronavirus HKU15, porcine epidemic diarrhoea virus, porcine astroviruses, Mamastrovirus 3, porcine torovirus and rotavirus C, and occasionally captured viruses absent from snout swabs at the same visit (Supplementary Fig. 8). Direct exposure to blood, saliva and other animal fluids was therefore a poor proxy for surveillance breadth, although the sample type retained value for selected viruses.

### Dominant viruses drive similarity while less common taxa reveal the blind spots

Sequentially removing each viral taxon and recalculating the difference in Jaccard distance showed that a small group of viruses accounted for much of the similarity between snout swabs and different environmental sample types. Coronavirus HKU15, porcine epidemic diarrhoea virus, porcine torovirus, enterovirus G and several astroviruses made the largest contributions (Fig. 2d; Supplementary Table 14). The same taxa dominated most pairwise comparisons (Supplementary Fig. 9; Supplementary Table 15) and together accounted for more than 80% relative abundance in each sample type (Fig. 2e). Similarity between sample types was therefore driven by a small set of widely detected viruses, whereas less frequently detected taxa accounted for the compositional differences between them.

Bray-Curtis dissimilarity, which accounts for viral read abundance, also separated sample types (PERMANOVA, FDR-adjusted P = 0.001, R² = 0.11; Supplementary Fig. 10a). Holding-pen swabs, faeces, wastewater, carcass wash water and ropes did not differ significantly from snout swabs in pairwise comparisons, whereas air and slaughter-box swabs remained distinct in virus composition (Supplementary Fig. 10a-c; Supplementary Tables 16-18). The same statistical separation therefore reflected opposite biological patterns: air contributed additional viral taxa beyond those detected in snout swabs, whereas slaughter-box swabs recovered only a restricted subset of the viral taxa present.

The blind spots were clearest among taxa absent from snout swabs. Mamastrovirus pig/HL/China was detected in both snout and environmental samples on one visit but only environmentally on six others. Porcine astrovirus 3 was shared on four visits and detected only environmentally on four more. Rotavirus A, porcine type C oncovirus, Aichivirus C and Sapporo virus also contributed strongly to dissimilarity between snout and environmental profiles (Supplementary Fig. 11). Several are associated with non-respiratory shedding, illustrating what is lost when surveillance is confined to one anatomical site.

Neither environmental-only nor snout-swab-only detections should be treated as mistakes by the other approach. Snout swabs represented selected animals and one anatomical compartment. Environmental sample types integrated material deposited through different routes and at different stages of slaughter. Concordance provided a useful benchmark, but the discordant detections contained much of the surveillance information.

### Environmental and pig-derived contigs show similar identities to shared reference sequences

Environmental contigs were compared with pig-derived contigs by aligning both against a custom database of viral proteins identified from snout-swab assemblies. Among all alignment pairs, 46,683, or 43.8%, differed by less than 1% in identity, representing 56 viral species (Supplementary Fig. 12; Supplementary Table 19). At a 3% threshold, 92,352 pairs, or 86.7%, representing 138 species showed similar identities to the shared reference. Coverage agreement was lower: 4,419 pairs, or 4.1%, differed by less than 1%, and 52,993, or 49.8%, differed by less than 3%.

When identity and coverage were considered together, 51,050 paired alignments, or 47.9%, representing 51 species differed by less than 3% for both measures. Coronavirus HKU15, mamastrovirus 3, rotavirus B and porcine astrovirus 2 fell within this range (Supplementary Table 19). Only 2.5% of alignment pairs differed by more than 10% in identity, although 19.6% differed by more than 10% in coverage. Porcine torovirus and porcine epidemic diarrhoea virus showed close identity agreement but larger differences in coverage, whereas rotavirus G and several bocaparvoviruses showed greater divergence across both measures.

### Combining environmental sample types gives consistent recovery across two slaughterhouses

Virus composition differed between Phnom Penh and Takeo (Jaccard PERMANOVA, P = 0.001, R² = 0.06; Fig. 3a). Observed richness was lower at Takeo (Fig. 3b), although Phnom Penh contributed six visits (72 samples) and Takeo three (31 samples), and the facilities differed in throughput and animal movement. Despite these ecological and sampling differences, the mean number of snout-swab detections recovered per visit by each environmental sample type did not differ significantly between locations, except that air contributed fewer sample-specific detections at Takeo (Supplementary Fig. 13a,b).

**Figure 3.**
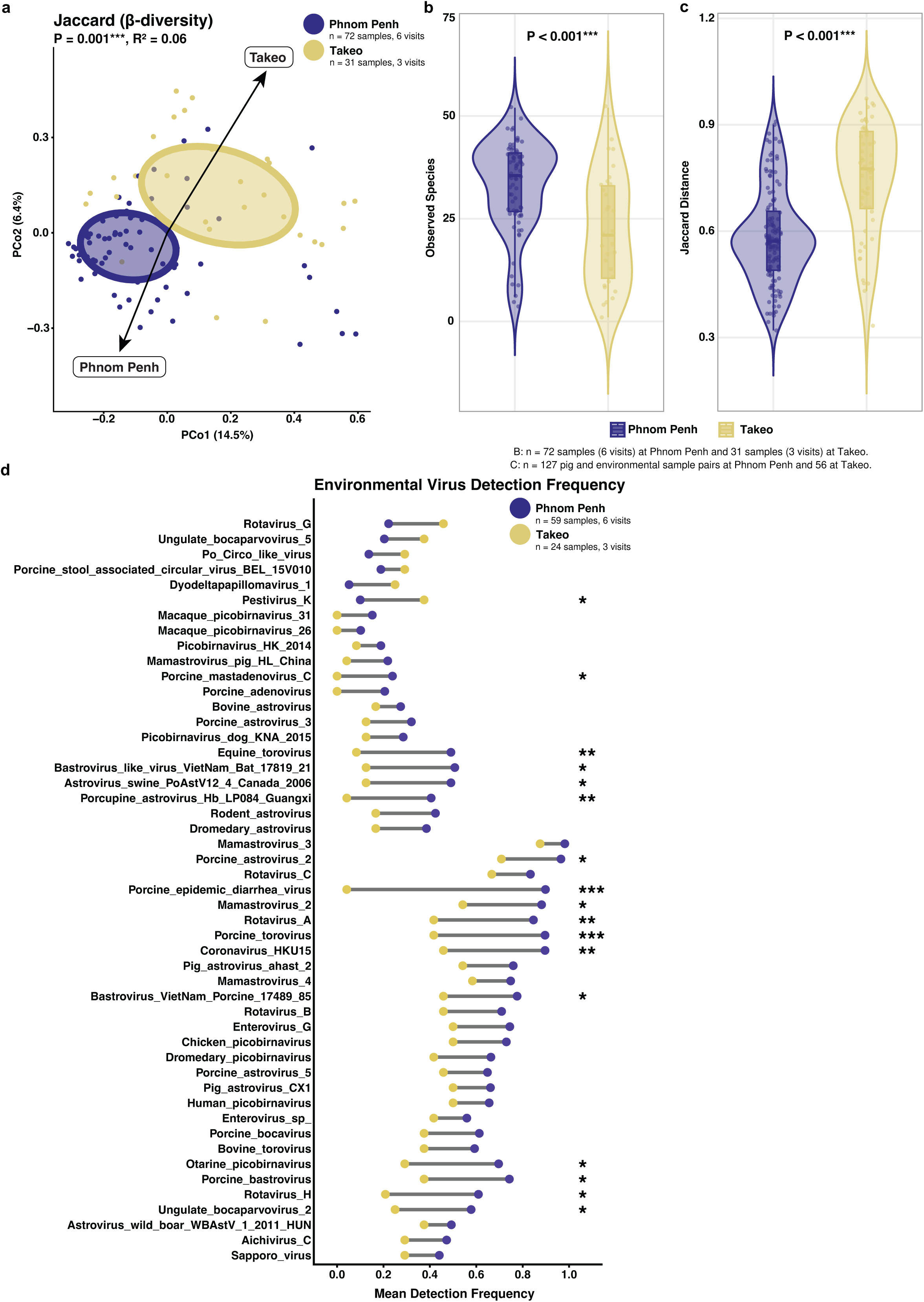
Virus composition and environmental detection frequency differ between slaughterhouses. a, Principal coordinate analysis of Jaccard dissimilarity at Phnom Penh (72 samples, 6 visits) and Takeo (31 samples, 3 visits). The arrow indicates the shift between site centroids. PERMANOVA tested differences in virus composition between sites. b, Observed viral richness by location (Phnom Penh, 72 samples, 6 visits, Takeo, 31 samples, 3 visits). c, Jaccard distance between pig snout swabs and environmental samples collected at the same visit (127 pig and environmental sample pairs at Phnom Penh and 56 at Takeo). d, Mean detection frequency by viral taxon and location, calculated as the mean across visits of the proportion of environmental samples in which each taxon was detected (Phnom Penh, 59 environmental samples, 6 visits, Takeo, 24 environmental samples, 3 visits). Wilcoxon rank-sum tests were used for two-group comparisons, and Fisher’s exact tests compared detection frequencies. P values were adjusted for false discovery using the Benjamini-Hochberg method. Adjusted P values are annotated as * P < 0.05, ** P < 0.01 and *** P < 0.001.

Individual environmental samples were more variable at Takeo and more distant from concurrently collected snout swabs than those from Phnom Penh (Fig. 3c). Sample types at Phnom Penh therefore resembled one another more closely, whereas at Takeo each sample type captured a different part of the virus composition. When detections were combined across sample types for each site and date, the distance to snout swabs was similar at both locations (Supplementary Fig. 14a,b), indicating that Takeo relied on combining a greater number of individual samples to achieve the same recovery. Coronavirus HKU15, porcine epidemic diarrhoea virus and rotavirus A were detected more frequently in Phnom Penh, whereas Pestivirus K was more frequent in Takeo (Fig. 3d; Supplementary Table 20). The usefulness of any single sample type therefore depended on the facility, whereas combining sample types gave consistent recovery at both.

### Environmental sampling reveals viral RNA associated with multiple hosts

Environmental samples detected 12 additional swine-associated viral taxa that were never identified in snout swabs (Fig. 4a). Porcine respirovirus 1 appeared in air, faeces and several other sample types across multiple dates. Air also detected Getah virus, Sapporo virus and an additional swine astrovirus on individual visits, whereas faeces uniquely detected porcine bocavirus 4 and porcine bufavirus. Astrovirus detections absent from snout swabs recurred across sample types and visits.

**Figure 4.**
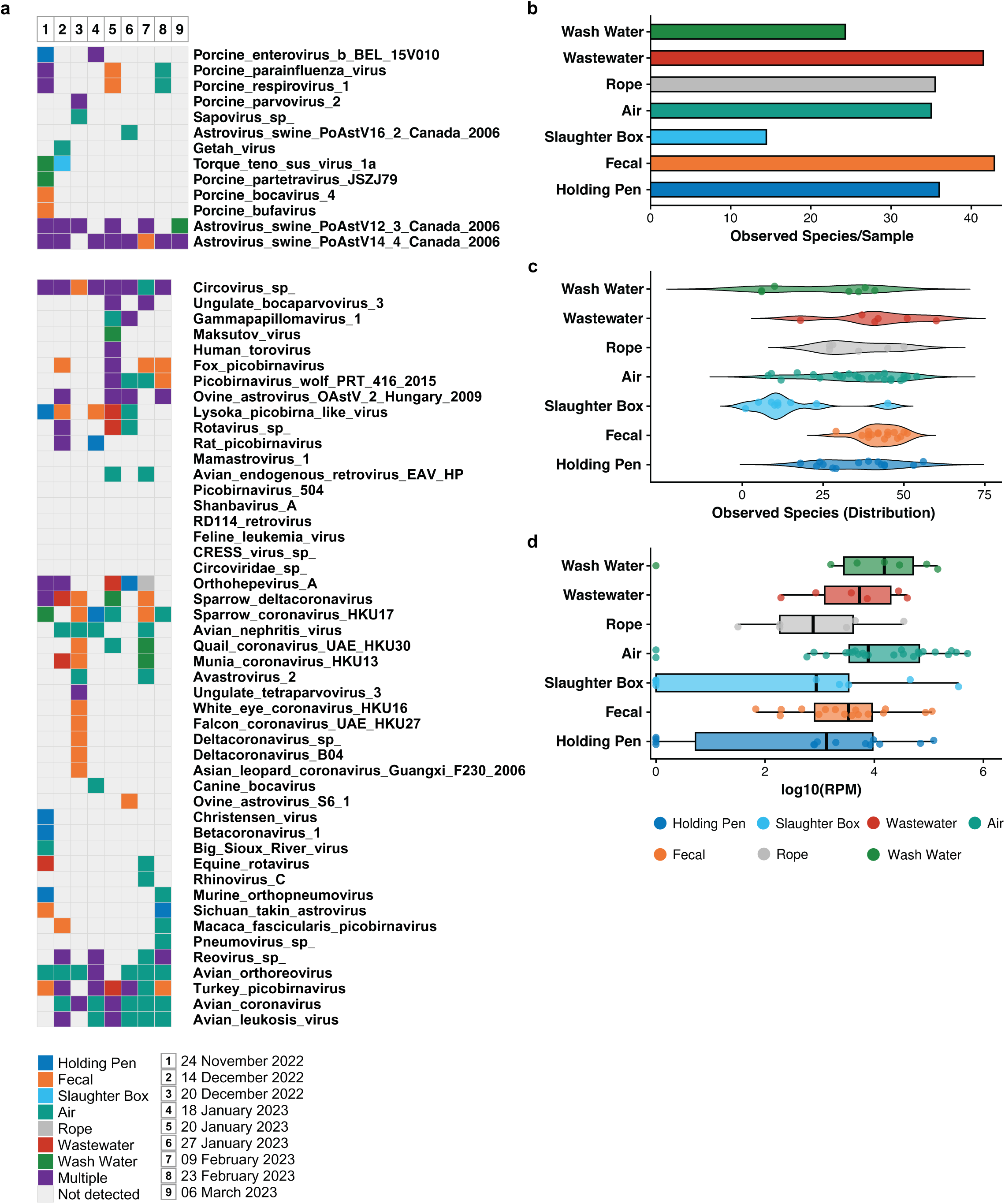
Environmental samples capture additional swine-associated and other host-associated viral taxa. a, Detection of viral taxa not identified in pig snout swabs during the study, by visit (1 to 9). Colours indicate the environmental sample type contributing each detection, and purple indicates detection in more than one sample type. Light grey indicates that the taxon was not detected in that visit and does not represent a diagnostically negative result. b, Mean number of additional taxa detected by each environmental sample type. c, Distribution of additional detections across individual samples. d, Abundance of additional taxa expressed as log10 reads per million. Host associations reflect taxonomic assignment and do not establish the source host, infection, viability or transmission.

Air, faeces, wastewater, holding-pen swabs and ropes each contributed more than 30 detections absent from snout swabs, with no significant differences among them (Fig. 4b-d; Supplementary Fig. 15a). Slaughter-box swabs contributed the fewest and significantly fewer than every other environmental sample type. The additional reach came from several imperfect samples rather than one universally superior sample type.

Environmental samples also contained viral detections associated with avian hosts, including members of the Astroviridae, Retroviridae, Reoviridae and Coronaviridae (Fig. 4a). Avian coronavirus, avian orthoreovirus and avian leukosis virus were detected during seven or eight of the nine visits and were most frequently recovered from air. Several human-associated detections were also made, including human coronavirus OC43, rhinovirus, Mamastrovirus 1 and human torovirus. These assignments identify viral nucleic acid associated with several host groups, not the source host, active infection, viability or transmission.

The multi-host detections were consistent with a facility that receives animals, people, vehicles, equipment, water and waste. Poultry slaughter was observed during one visit, and pigs commonly originate from smallholder systems in which several species may be raised together. Wildlife and neighbouring premises could also contribute material. These sources could not be resolved.

## Discussion

Environmental sampling in a slaughterhouse can be used to survey the virus diversity of the pigs processed there, offering a more efficient and cost-effective alternative to sampling individual animals. In this study, combining multiple environmental sample types recovered 96.6% of snout-swab viral taxa overall, exceeding 90.9% at every visit, and detected additional taxa associated with swine and other hosts. Air provided the widest view from one platform, but faeces, ropes, water and surfaces filled different blind spots in surveillance coverage. The slaughter box supplied a useful example of failure: repeated contact with animal fluids did not translate into broad viral recovery. Environmental metagenomics therefore complemented snout-swab sampling, capturing the same core virus diversity while extending detection to the wider slaughterhouse environment.

Most environmental surveillance programmes in pig-production systems have targeted individual viruses by PCR or RT-qPCR [33–37], including influenza A virus [38–41] and porcine epidemic diarrhoea virus [42, 43]. However, metagenomic studies of wastewater, air, farms and slaughterhouses can detect a broader range of viruses present but remain less common [44–47]. In addition, few studies have compared several environmental sample types with samples from the pigs themselves. It has therefore been unclear how well environmental samples represent the viruses present in pigs. By comparing seven environmental sample types with pig snout swabs collected at the same visits, this study measured how much of the pig virus diversity each environmental sample type recovered.

Each sample type recovered a different set of viruses, reflecting the material it collects. Wastewater and faeces accumulate enteric viruses [44, 45], air collects respiratory secretions, dust and resuspended material [38–40, 46], and ropes and holding-pen surfaces are repeatedly contacted by oral, nasal and skin secretions. Because each sample type reflects a different route of viral release, no single sample type captured every taxon, and some taxa found in snout swabs were absent from the environmental samples. Some of these absences are consistent with the shedding routes of the viruses concerned. Pestiviruses, for example, can be transmitted congenitally and maintained by persistently infected animals that shed through serum and other secretions [48–50], and the transmission route of porcine circovirus 5 remains uncertain [51]. In contrast, the failure to recover porcine mastadenovirus A and porcine stool-associated circular virus 5, which are associated with faecal shedding [52, 53], could reflect low abundance, intermittent shedding or incomplete sampling.

Air provided the widest environmental view from a single sample type. Each air sampler collected material released by multiple animals and activities over three hours, whereas a swab represents one animal, surface or shedding route. Air detections cannot be attributed to individual animals, cannot distinguish active shedding from resuspended material, and do not establish infection or viability. Air sampling is therefore best suited to facility-level surveillance, where it can show which viruses are present and guide targeted investigation. Air sampling has been used in the same way in live-poultry markets [27], dairy farms [54], mink farms [55], camel barns [56] and commercial swine facilities [38, 40, 57].

On the other hand, the slaughter box was visibly and repeatedly exposed to blood, saliva and other animal material, yet its swabs recovered the fewest taxa. This may reflect its additional use for de-hairing carcasses with hot or boiling water, which could wash away material and degrade viral nucleic acid on the surface. Contamination that looks obvious is not necessarily surveillance that works. Even so, slaughter-box swabs repeatedly detected coronavirus HKU15 and porcine epidemic diarrhoea virus and occasionally captured detections absent from snout swabs. This indicates that slaughter-box swabs are better suited to targeted detection of specific viruses than to broad surveillance of virus diversity. Sample selection should therefore be guided by the surveillance objective and not by the degree of visible contamination.

Virus richness and the overlap in detections among sample types differed between the two slaughterhouses. Phnom Penh had greater richness and more taxa shared among sample types, which may reflect higher throughput of animals. In Takeo, virus composition varied more among sample types, so recovery depended on combining a greater number of samples. Ventilation, cleaning, disinfection and facility design may also influence recovery from air and surfaces [58, 59], but these factors were not measured. The comparison supports the consistency of combining sample types across two contrasting facilities.

The avian and human-associated detections widened the view and narrowed the certainty. Poultry slaughter was observed once, yet avian-associated detections appeared repeatedly, especially in air. Possible sources included adjacent premises, transport, equipment, workers, wild birds, water and feed. Similar multi-host detections have been identified through environmental metagenomics in Cambodian live-poultry markets [27]. These findings show that the environment integrates material associated with several hosts. However, this does not allow for identification of the source host or demonstrate active infection, viability or transmission.

Collecting every sample type gave the widest coverage of viral taxa, but routine surveillance is limited by time, sequencing capacity and cost. Future work should therefore identify the smallest combination of sample types that retains useful detection across facilities and seasons. In these data, air provided the widest coverage, with faeces, ropes, holding-pen swabs and water filling different gaps in detection. Future work comparing the effort, cost and turnaround time of each sample type across facilities and seasons could define the most efficient combination for routine use. Unexpected or epidemiologically important metagenomic detections should be confirmed by targeted testing and followed by investigation of the animals, since detection alone does not establish infection.

Several limitations should be considered when interpreting these results. The study included two slaughterhouses and nine visits, with unequal sampling between locations. Recovery was measured against snout swabs, which sample the upper respiratory tract and oral cavity and may not represent viruses at other body sites. No influenza A virus was detected in snout swabs or environmental samples, although the capture panel covers influenza A viruses and the same approach has previously detected avian influenza viruses in air and environmental samples from live poultry markets in Cambodia [27], so panel design does not explain the absence. Influenza A virus has previously been identified through sequencing and serological studies in pigs at other slaughterhouses in Cambodia not sampled in this study, although detection frequency in Cambodian pigs is reportedly low [60–62]. Environmental sample types also differed in how they were collected, with each air sample taken by a single air sampler and all other samples made up of multiple swabs pooled together, which may have influenced the number of taxa recovered from each.

These findings support a tiered approach in which air sampling provides broad coverage, other environmental sample types fill gaps in detection, and sampling of animals is used to identify the host of unexpected detections. Overall, this evidence suggests that environmental metagenomics offers a scalable and resource-efficient means of enhancing surveillance for emerging viruses, particularly in settings where conventional animal-based sampling is impractical. Broader implementation of this approach at high-risk human-animal interfaces globally could strengthen early warning systems and improve pandemic preparedness. More fundamentally, it offers a practical route to operationalising One Health surveillance, providing a single, tractable workflow capable of capturing viral threats across the human, animal and environmental interface.

## Contributors

P.C. performed the bioinformatic analysis, curated and validated the data, produced the visualisations and wrote the manuscript. V.H. and J.Y.S. curated the data and reviewed and edited the manuscript. S.To., S.Si., B.Si., V.O., S.N. and K.C. collected the samples and prepared them for next-generation sequencing. J.N., V.C., B.Se., M.H., S.Sa., S.Tu. and F.C. contributed to conceptualising the study. C.F., Y.C.F.S., G.J.D.S. and E.A.K. conceived the study and reviewed and edited the manuscript. All authors contributed substantially to the discussion of the content.

## Acknowledgements

The investigators thank everyone involved in the critical discussions and review of this manuscript, as well as all those contributing to virus surveillance and response in the Kingdom of Cambodia. This includes teams at the National Institute for Public Health, the Cambodian Communicable Disease Control Department, Ministry of Health, the National Animal Health and Production Institute, General Directorate of Animal Health and Production, Ministry of Agriculture, Forestry, and Fisheries, the World Health Organization, the United States Centers for Disease Control and Prevention, provincial health and agriculture directors, slaughterhouse managers, middlemen, vendors, and sellers, and the Virology Unit at Institut Pasteur du Cambodge, all of whom contributed to this study. Research reported in this publication was supported by the Bill and Melinda Gates Foundation (INV-049293), the National Institute of Allergy and Infectious Diseases of the National Institutes of Health Centers for Research in Emerging Infectious Diseases under subaward #12-312-0217530 issued under prime award #1U01AI151378-01, the Food and Agriculture Organization of the United Nations (FAO), contract 75N93021C00016 from the National Institute of Allergy and Infectious Diseases, NIH, Department of Health and Human Services, USA, and by the Duke-NUS Signature Research Programme funded by the Ministry of Health, Singapore. P.C. is supported by the National Research Foundation, Singapore under its Postdoctoral Fellowship (Postdoctoral Award FY2025-NRF-PA-0002). The text as published does not necessarily represent the official views of the US-CDC, WHO, FAO, or the National Institutes of Health. The content is solely the responsibility of the authors.

## Ethics statement

All research complied with relevant ethical regulations. Sample collection protocols were approved by the National Ethics Committee for Health Research, Ministry of Health, Cambodia (approval numbers NECHR013, NECHR143, NECHR149 and NECHR320; approved in 2021-2022). Cambodia does not have a dedicated animal ethics review board. Animal sampling was conducted in collaboration with the National Animal Health and Production Institute and with authorization from the General Directorate of Animal Health and Production, Ministry of Agriculture, Forestry and Fisheries.

