## Supplemental Figure 1 for "Air and environmental sampling expand metagenomic viral surveillance in Cambodian pig slaughterhouses"

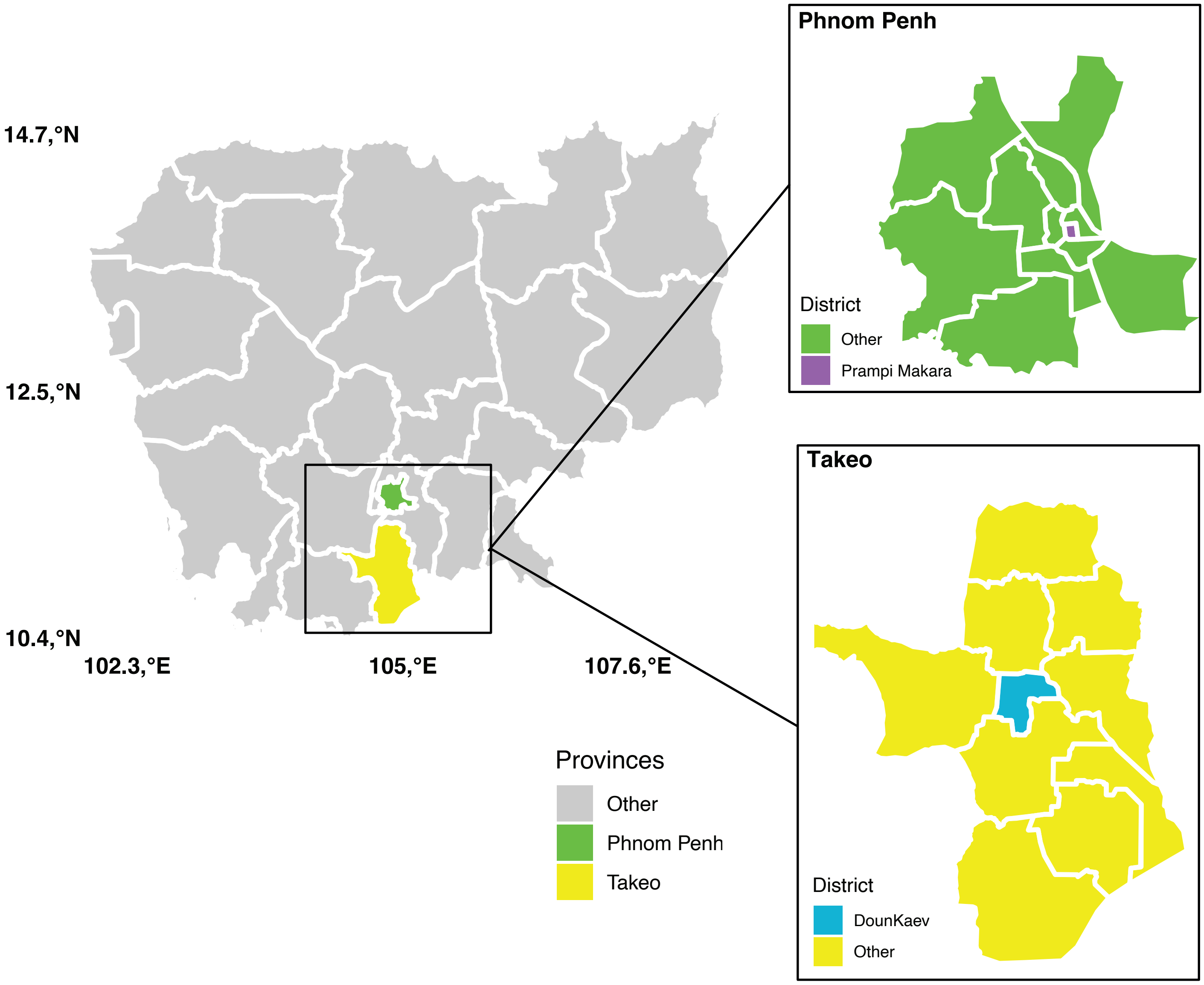


**Supplementary Figure 1: Geographic location of pig slaughterhouse sampling locations used in Cambodia.** The main panel presents a map of Cambodia highlighting the two provinces in which surveillance was conducted, Phnom Penh and Takeo. Insets show enlarged views of each province, indicating the specific districts where the pig slaughterhouses were located. In Phnom Penh, the slaughterhouse is located within the central district of Prampi Makara, whereas the Takeo slaughterhouse was situated in Doun Kaev district.
