## Supplemental Figure 2 for "Air and environmental sampling expand metagenomic viral surveillance in Cambodian pig slaughterhouses"

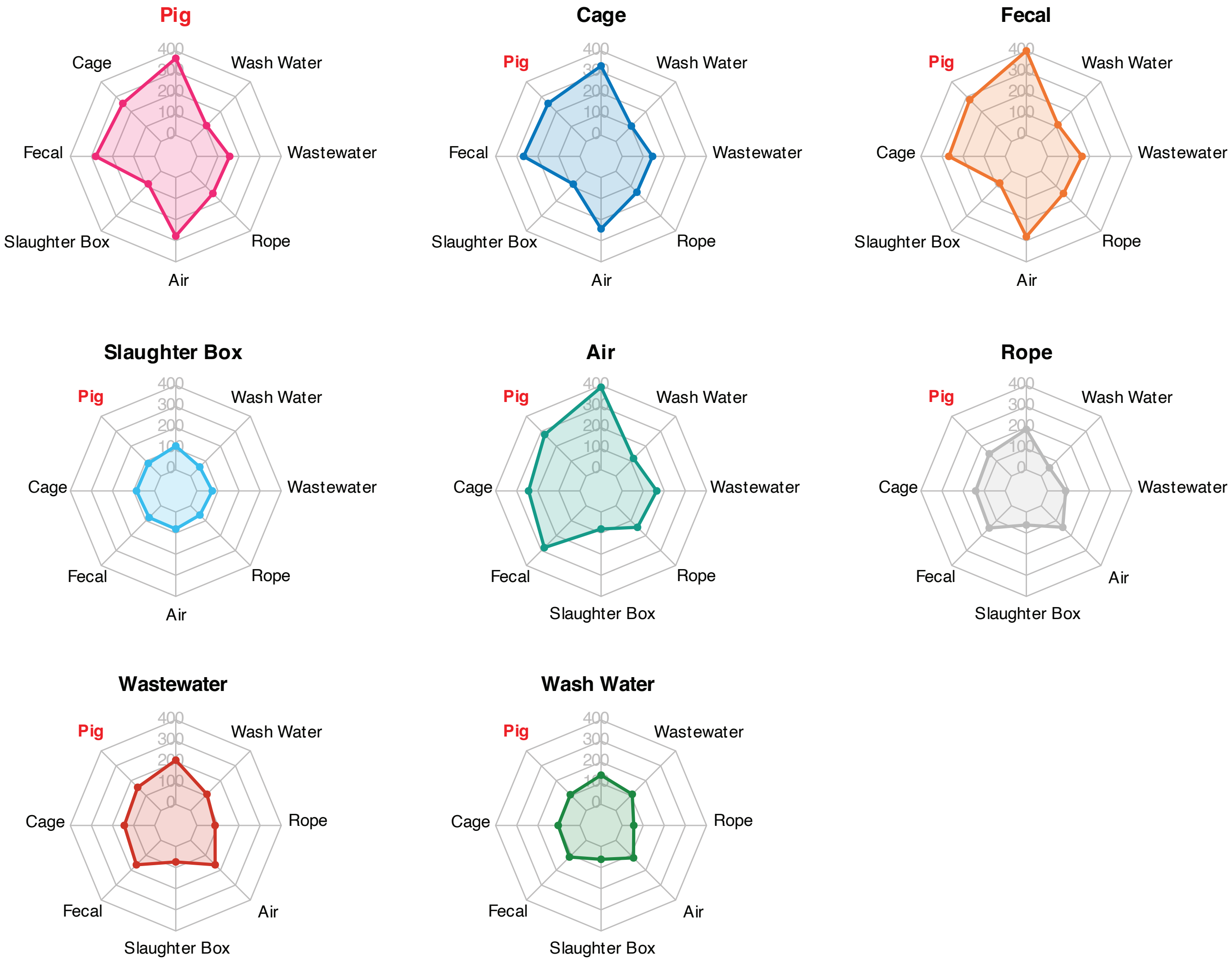


**Supplementary Figure 2: Comparison of virus detection frequency between pig and environmental samples.** The total number of unique virus detections (defined as the presence of a given virus in one sample type at a single time point) shared between different sample types is shown. Each radar plot is centered on a focal sample type, and the values along each axis indicate the number of detections shared between that focal sample type and the corresponding comparison sample type. Virus detections shared with pig samples are highlighted in red.
