## Supplemental Figure 3 for "Air and environmental sampling expand metagenomic viral surveillance in Cambodian pig slaughterhouses"

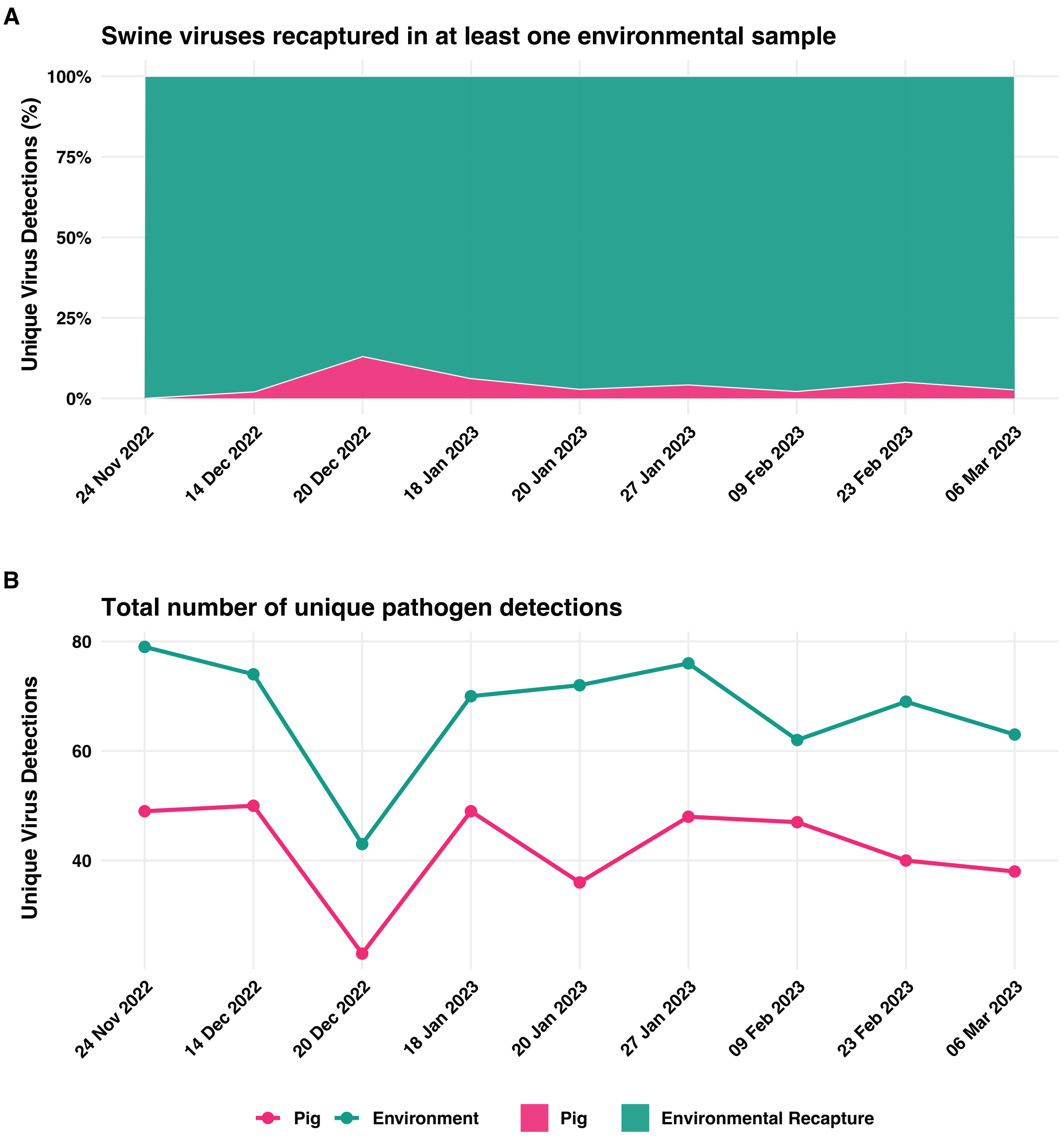


**Supplementary Figure 3: Unique virus detections in pig and environmental samples over time.** A) The percentage of swine virus detections identified in pig samples that were recaptured in at least one environmental sample was calculated for each sampling date. B) The total number of unique virus detections identified in pig and environmental samples was compared across sampling dates. Unique virus detections were defined as distinct viruses detected at least once within a sample group on a given sampling date, regardless of how many individual samples were positive.
