## Supplemental Figure 4 for "Air and environmental sampling expand metagenomic viral surveillance in Cambodian pig slaughterhouses"

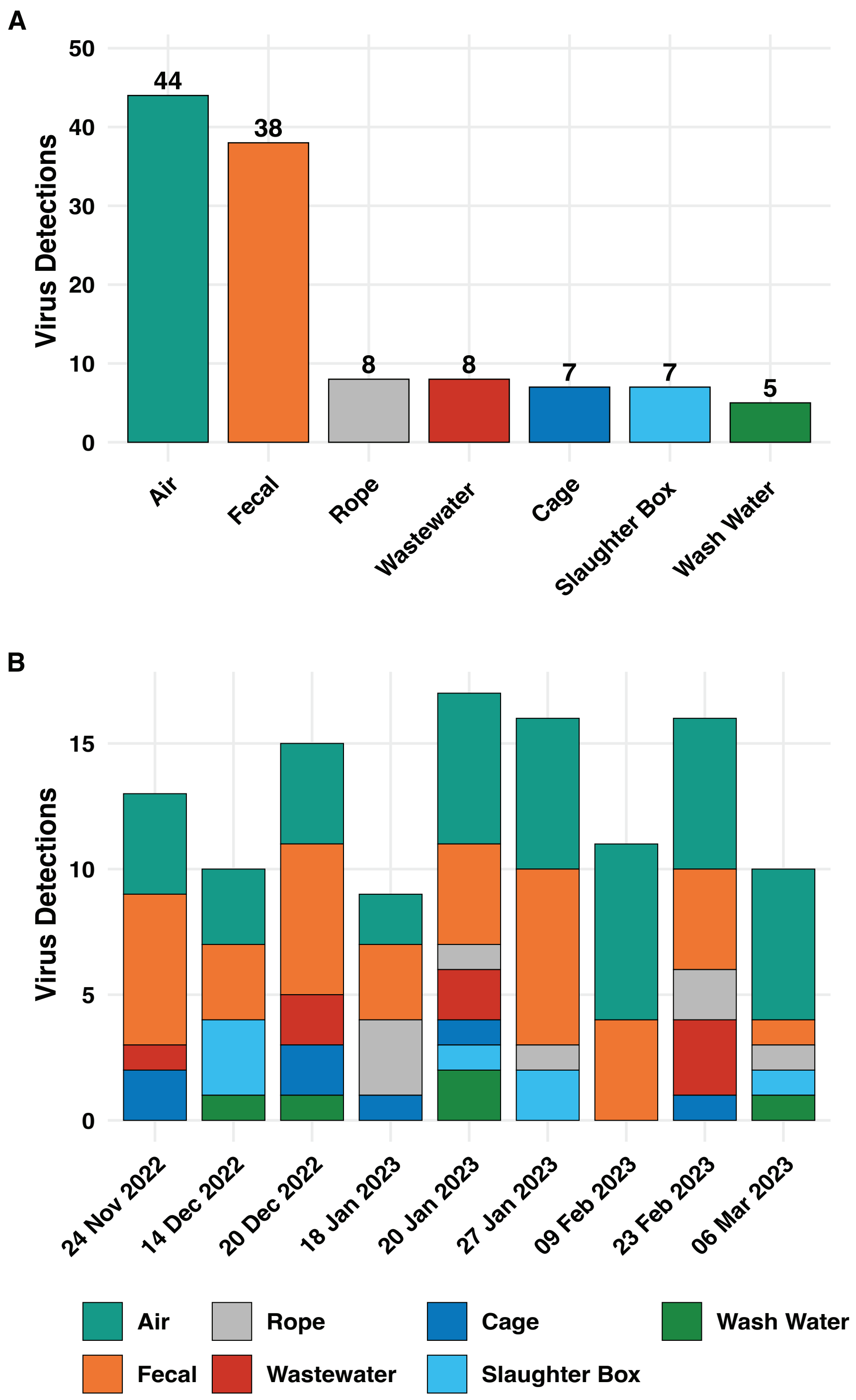


**Supplementary Figure 4: Viruses uniquely captured in the environmental samples.** A) The total number of unique virus detections identified exclusively within each environmental sample type is shown. B) The same exclusive detections are shown by sampling date, with stacked bars indicating the contribution of each environmental sample type at each time point. Unique detections were defined as distinct viruses detected at least once within a sample type on a given sampling date, regardless of how many individual samples were positive. Only detections found in one environmental sample type and not detected in pig or any other sample type on the same sampling date were included.
