## Supplemental Figure 5 for "Air and environmental sampling expand metagenomic viral surveillance in Cambodian pig slaughterhouses"

A

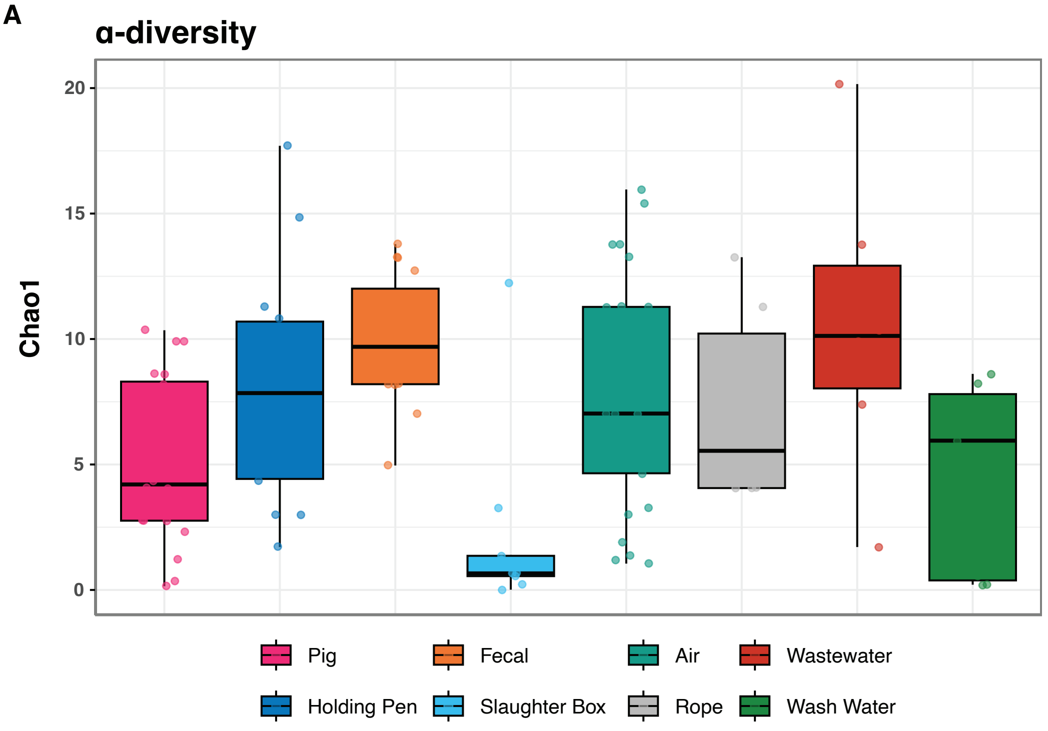


B

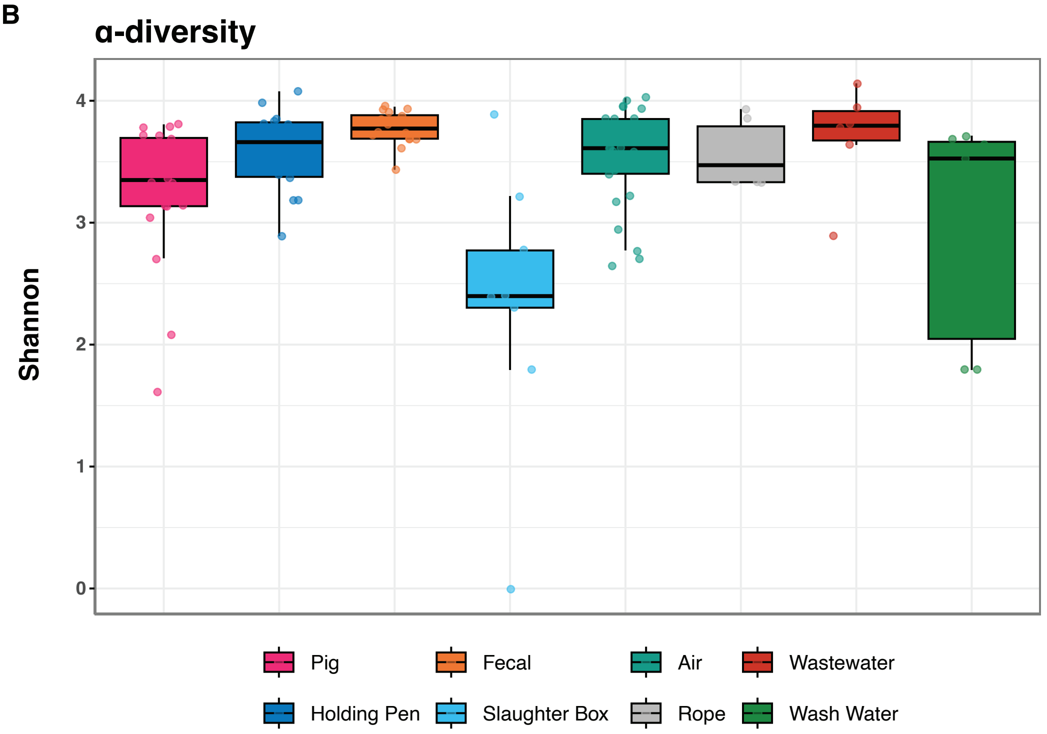


**Supplementary Figure 5: Environmental samples consistently capture the majority of swine virus pathogens.** α-diversity measures A) Chao1 and B) Shannon Index were calculated to compare the virus richness between different environmental sample types and the routine pig snout swab sampling method. Statistics were calculated using a Kruskal–Wallis with Dunn's post-hoc test. All P-values obtained were corrected for false discovery rate (FDR) using the Benjamini–Hochberg method. P-values for all pairwise comparisons can be found in Supplementary Tables 3-5.
