## Supplemental Figure 6 for "Air and environmental sampling expand metagenomic viral surveillance in Cambodian pig slaughterhouses"

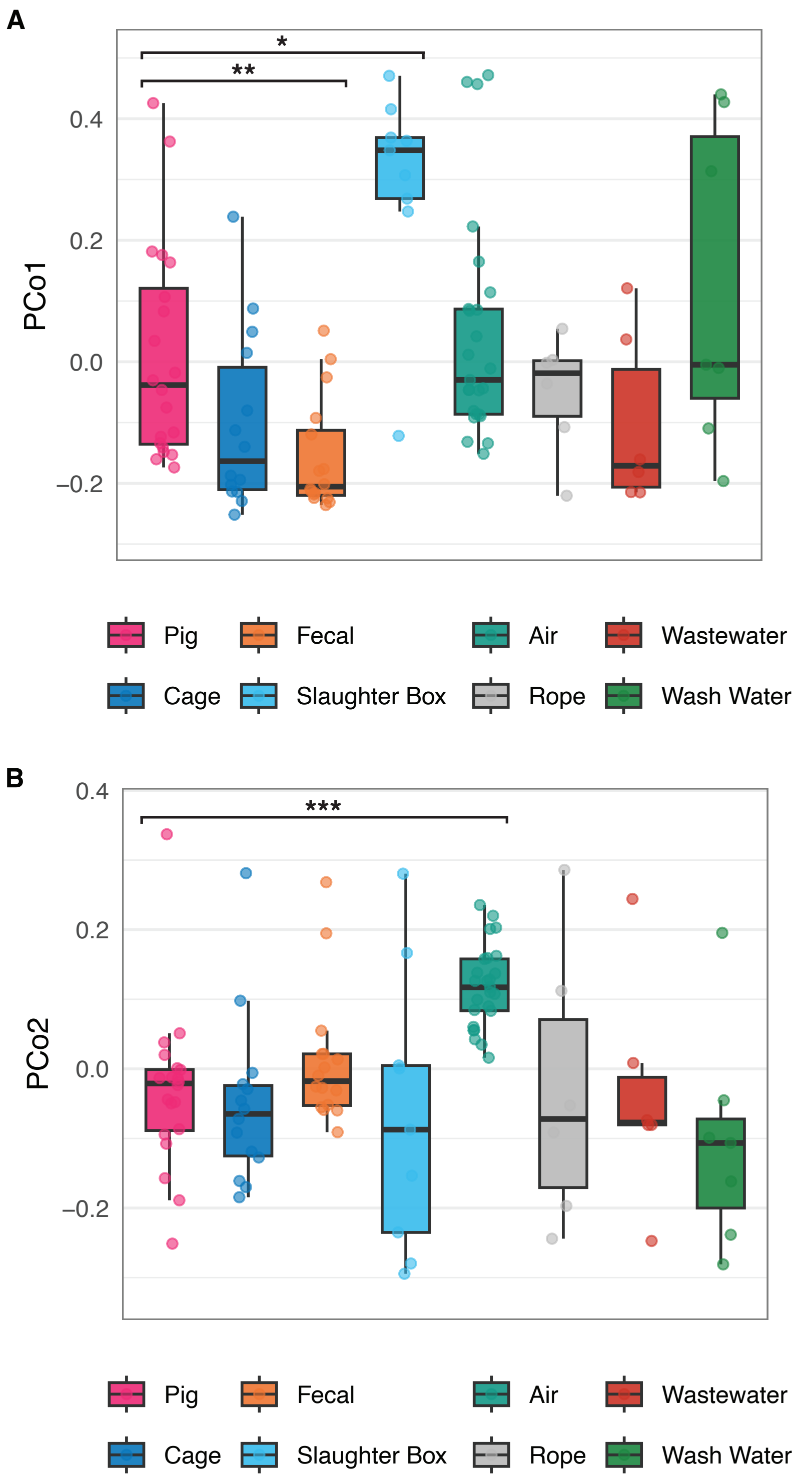


**Supplementary Figure 6: First and second principal coordinates derived from the Jaccard distance between samples shows significant virome separation.** The distribution of values along each coordinate axis A) PCo1 and B) PCo2 respectively. Statistics were calculated using a Kruskal–Wallis with Dunn's post-hoc test. All P-values obtained were corrected for false discovery rate (FDR) using the Benjamini–Hochberg method. P-values for all pairwise comparisons can be found in Supplementary Tables 7-8.
