## Supplemental Figure 7 for "Air and environmental sampling expand metagenomic viral surveillance in Cambodian pig slaughterhouses"

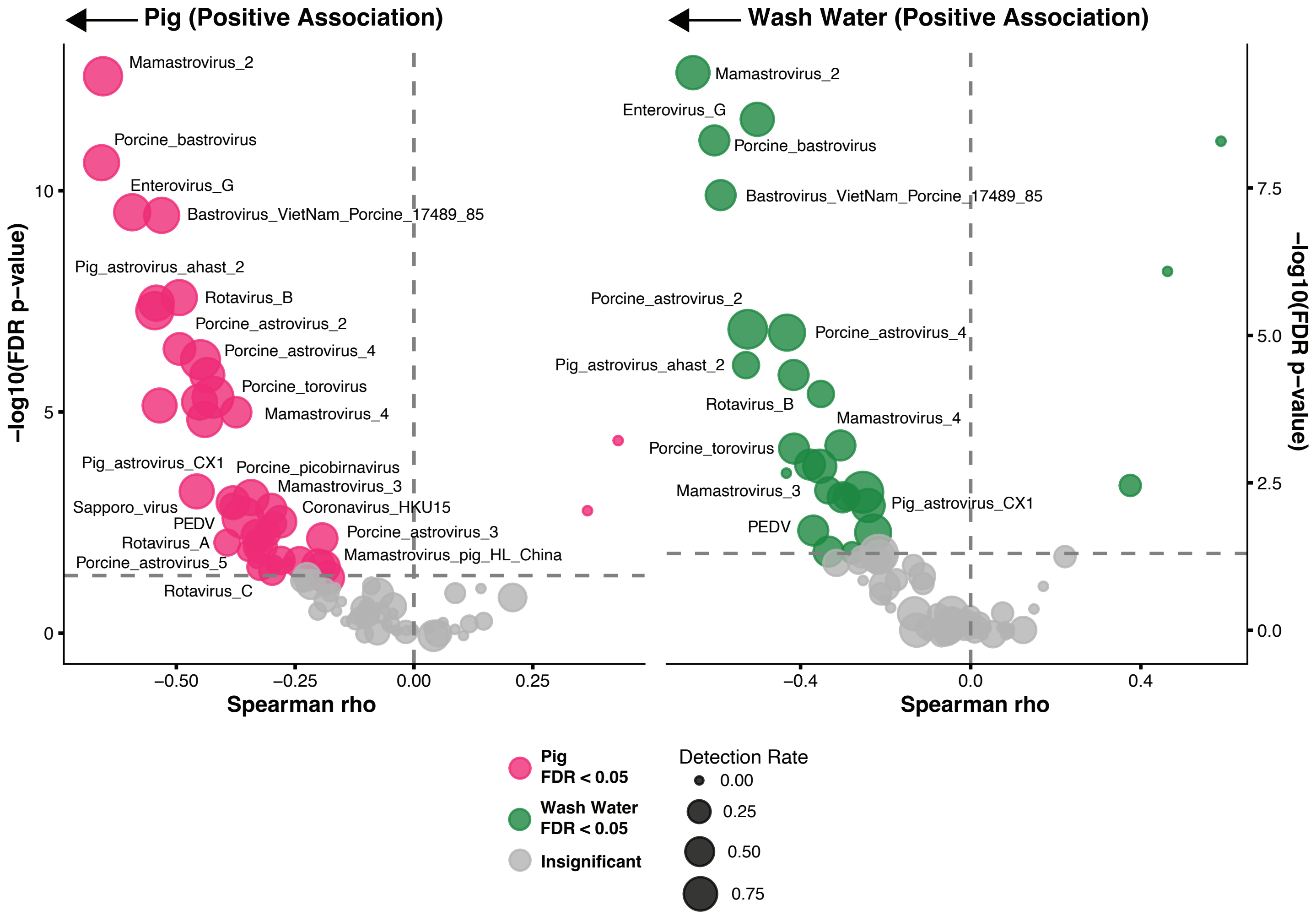


**Supplementary Figure 7: Swine viruses positively associate with pig snout swabs and the ES carcass wash water.** Spearman correlations between virus detection frequency and the distance to the median PCo1 and PCo2 coordinates of Jaccard β-diversity PCoA. The full list of significant taxa and corresponding correlation values can be found in Supplementary Tables 11 and 12.
