## Supplemental Figure 8 for "Air and environmental sampling expand metagenomic viral surveillance in Cambodian pig slaughterhouses"

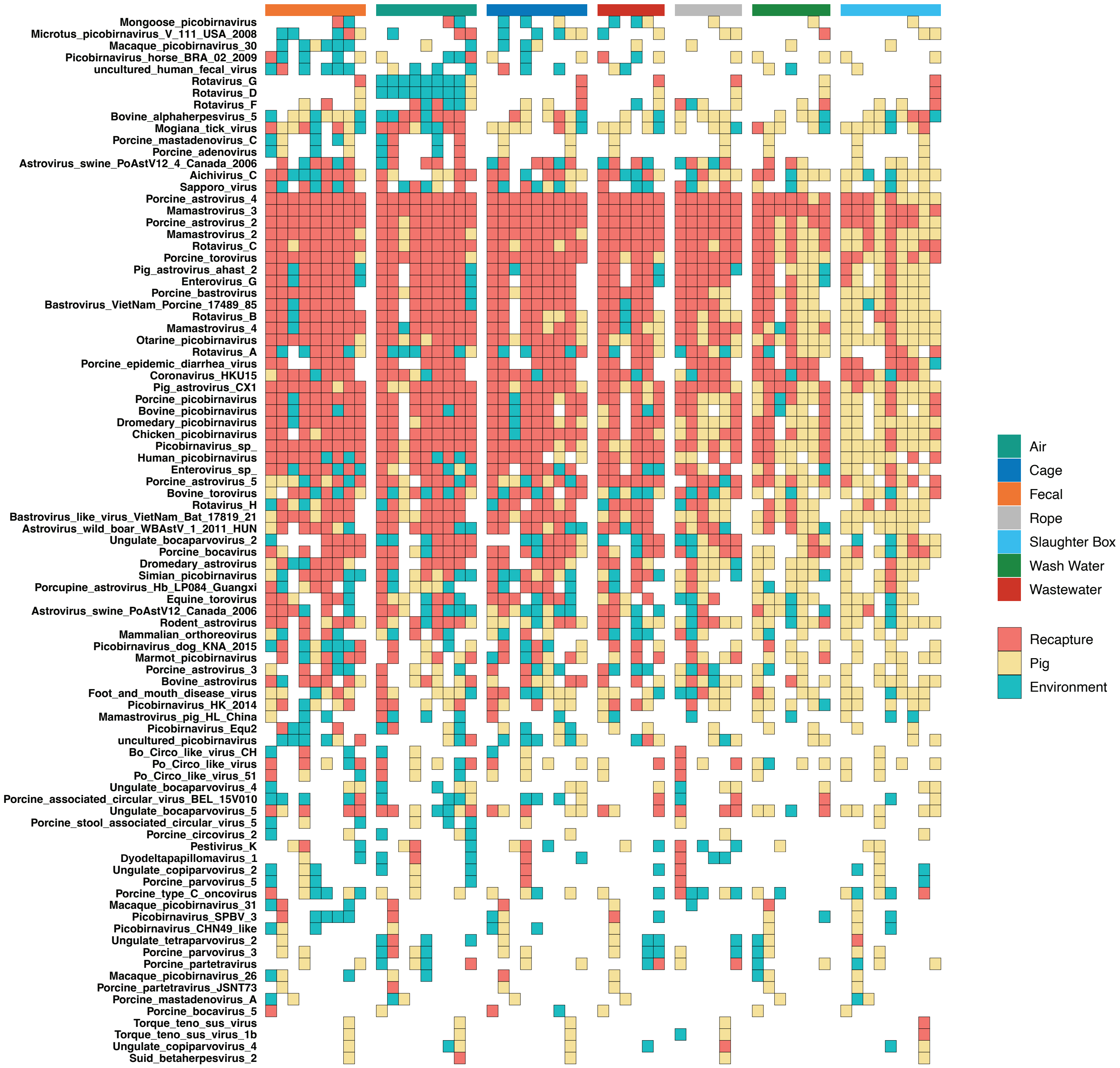


**Supplementary Figure 8: Detection of swine viruses through environmental sampling.** Heatmap showing 88 different virus species which were detected at least once in pig swabs over the course of the study. The red color shows when a virus was identified in pigs and at the exact same timepoint was recaptured in the environment. The pale yellow colour shows when a virus species was identified in pigs but failed to be recaptured through environmental sampling. The blue colour shows virus species which were found in the environment but failed to be detected in pigs. Rows of the heatmap represent virus species and columns represent individual environmental samples (annotated at the top of the heatmap).
