## Supplemental Figure 9 for "Air and environmental sampling expand metagenomic viral surveillance in Cambodian pig slaughterhouses"

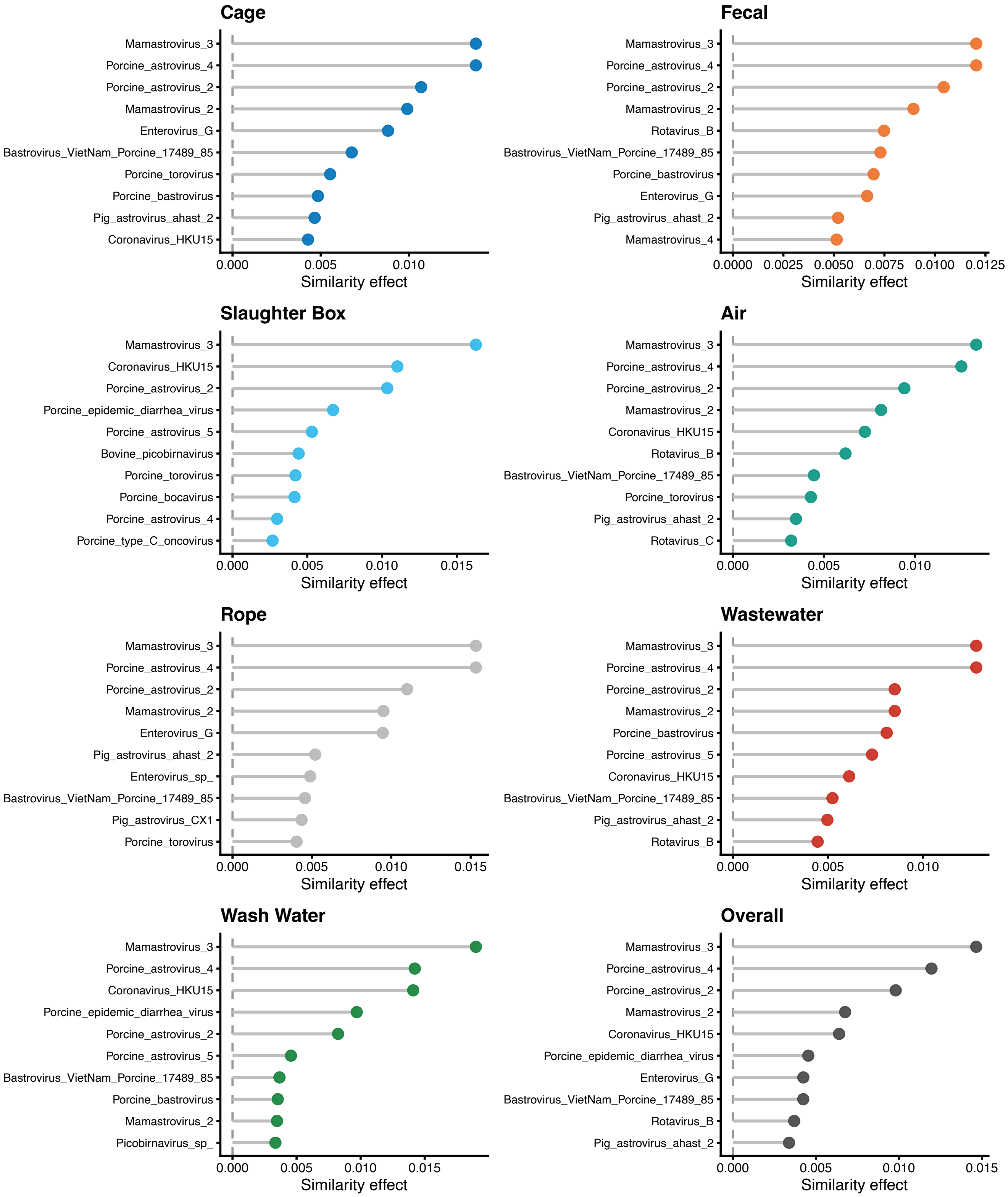


**Supplementary Figure 9: Top 10 viral taxa with the greatest contribution to compositional similarity between pig swabs and environmental samples.** Similarity effect was calculated by sequentially removing each virus from Jaccard β-diversity matrix and calculating the difference in distance. Supplementary Tables 14 and 15 provide the full set of values.
