## Supplemental Figure 10 for "Air and environmental sampling expand metagenomic viral surveillance in Cambodian pig slaughterhouses"

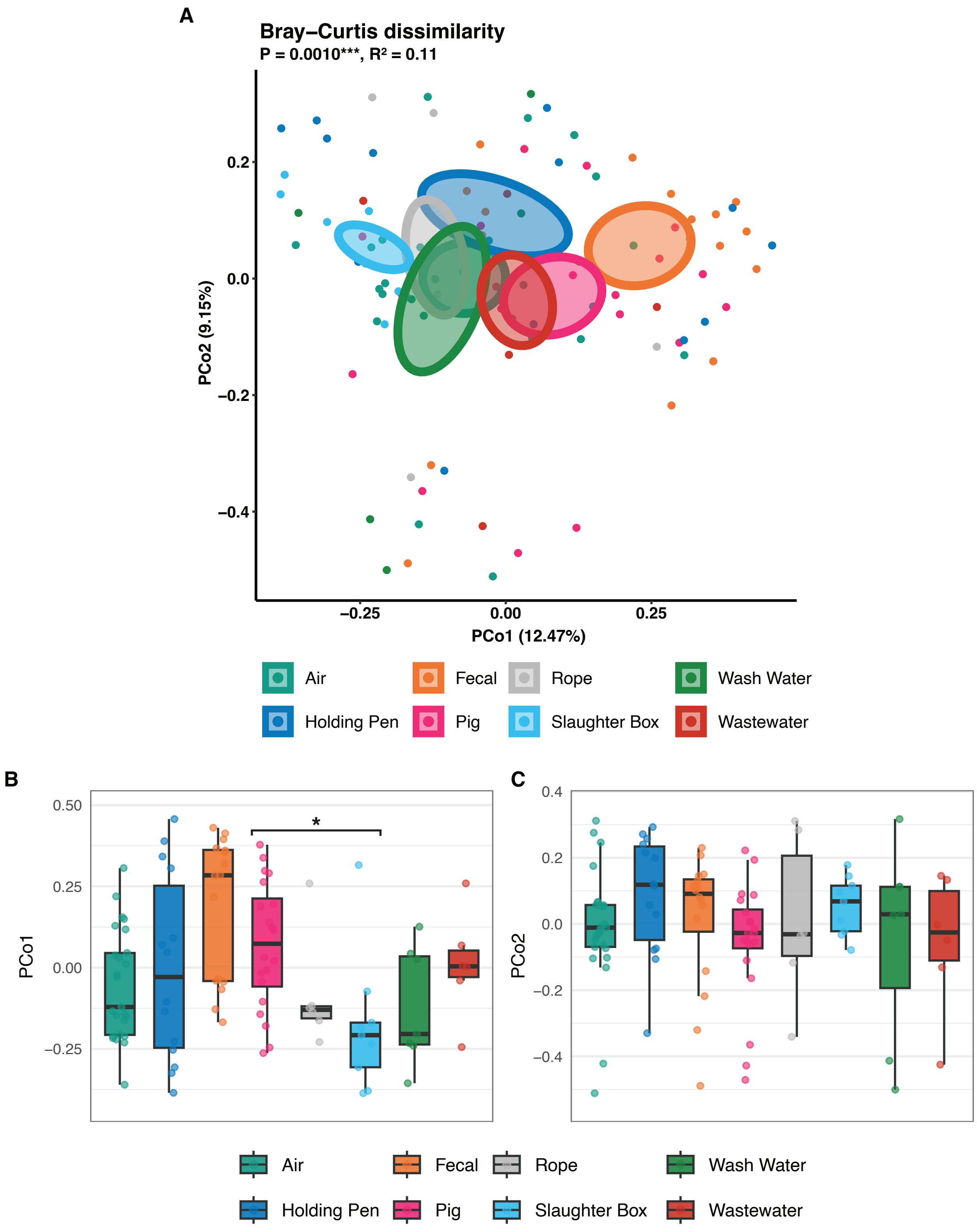


**Supplementary Figure 10: Bray-Curtis dissimilarity which accounts for virus abundance shows significant separation between sample types.** A) PCoA based on Bray–Curtis dissimilarity beta-diversity. Statistical significance of overall differences in virus composition between sample types was assessed by PERMANOVA. The distribution of values along each coordinate axis B) PCo1 and C) PCo2 respectively. Statistics were calculated using a Kruskal–Wallis with Dunn's post-hoc test. All P-values obtained were corrected for false discovery rate (FDR) using the Benjamini–Hochberg method. P-values for all pairwise comparisons can be found in Supplementary Tables 16-18.
