## Supplemental Figure 11 for "Air and environmental sampling expand metagenomic viral surveillance in Cambodian pig slaughterhouses"

**a**

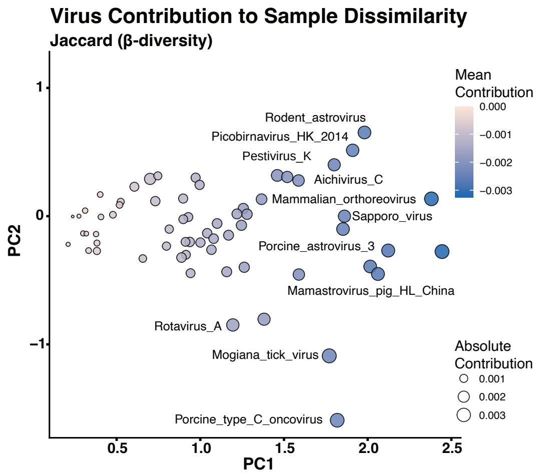


**b**

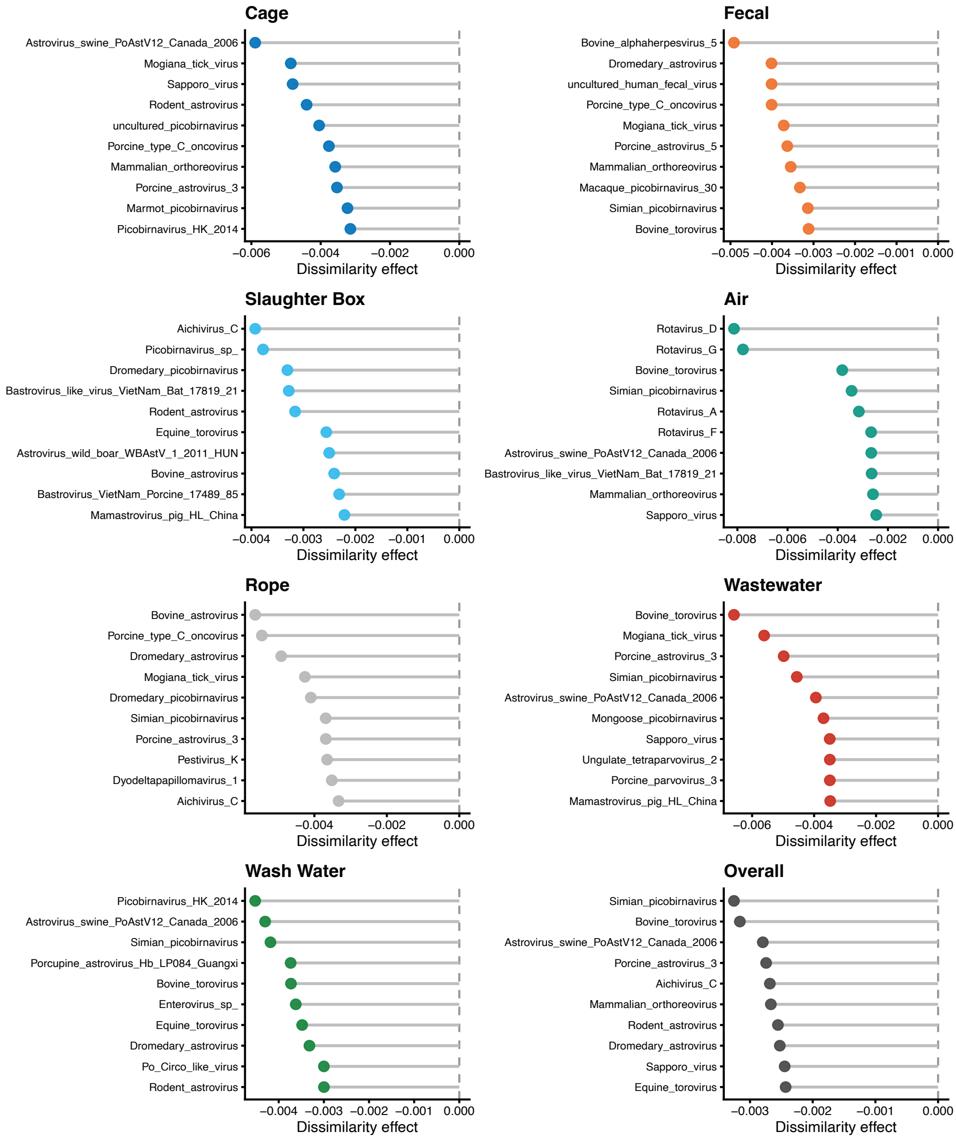


**Supplementary Figure 11: Taxa with the greatest contribution to compositional dissimilarity between pig swabs and environmental samples.** a, Principal coordinates analysis of the contribution of individual taxa to dissimilarity between pig and environmental samples, estimated by sequential removal of each taxon and recalculation of the Jaccard distance. Point colour indicates the mean contribution, and point size indicates the absolute contribution. b, Top 10 viral taxa with the greatest contribution to compositional dissimilarity for each sample type. Similarity effect was calculated by sequentially removing each virus from Jaccard β-diversity matrix and calculating the difference in the distance. Supplementary Tables 14 and 15 provide the full set of values.
