## Supplemental Figure 12 for "Air and environmental sampling expand metagenomic viral surveillance in Cambodian pig slaughterhouses"

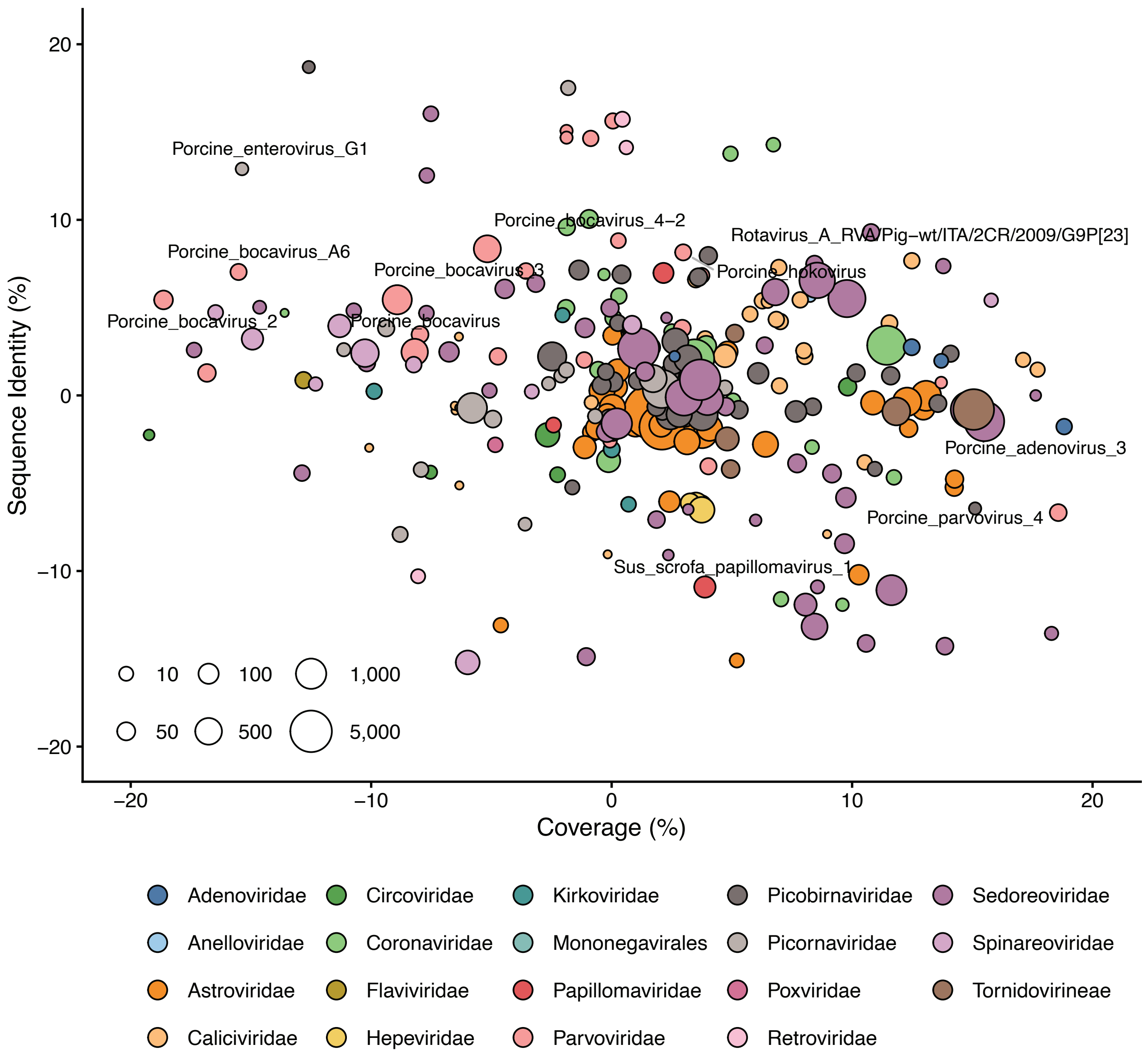


**Supplementary Figure 12: Majority of swine pathogens exhibit few differences in sequence similarity and coverage between pig and the environment.** Contig assemblies from the environment and from pigs were aligned to the same viral protein where the differences in coverage (%) and sequence identity (%) was calculated. Node size represents the number of contig pairs for a given species. Nodes are colored according to viral family.
