## Supplemental Figure 13 for "Air and environmental sampling expand metagenomic viral surveillance in Cambodian pig slaughterhouses"

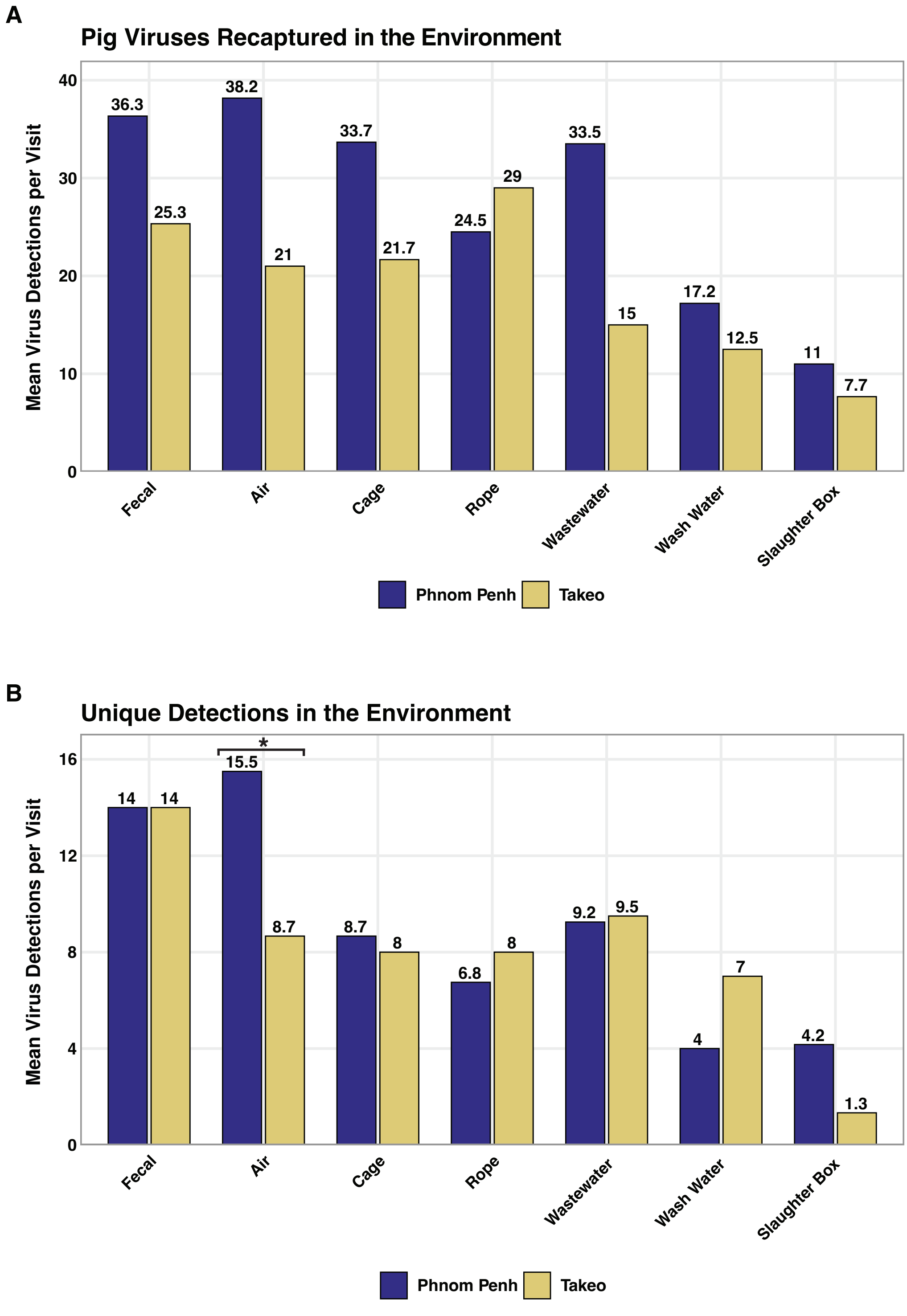


**Supplementary Figure 13: Number of virus species detected at different locations across each different environmental sample types.** A) The number of pig viruses recaptured in a given environmental sample type was calculated at each visit and averaged across all visits to each location. B) Using the same approach as A), where only unique virus detections were counted, where a virus was identified in an environmental sample but could not be detected in the pig samples collected at that visit. Phnom Penh, n = 6 visits and Takeo, n = 3 visits.
