## Supplemental Figure 14 for "Air and environmental sampling expand metagenomic viral surveillance in Cambodian pig slaughterhouses"

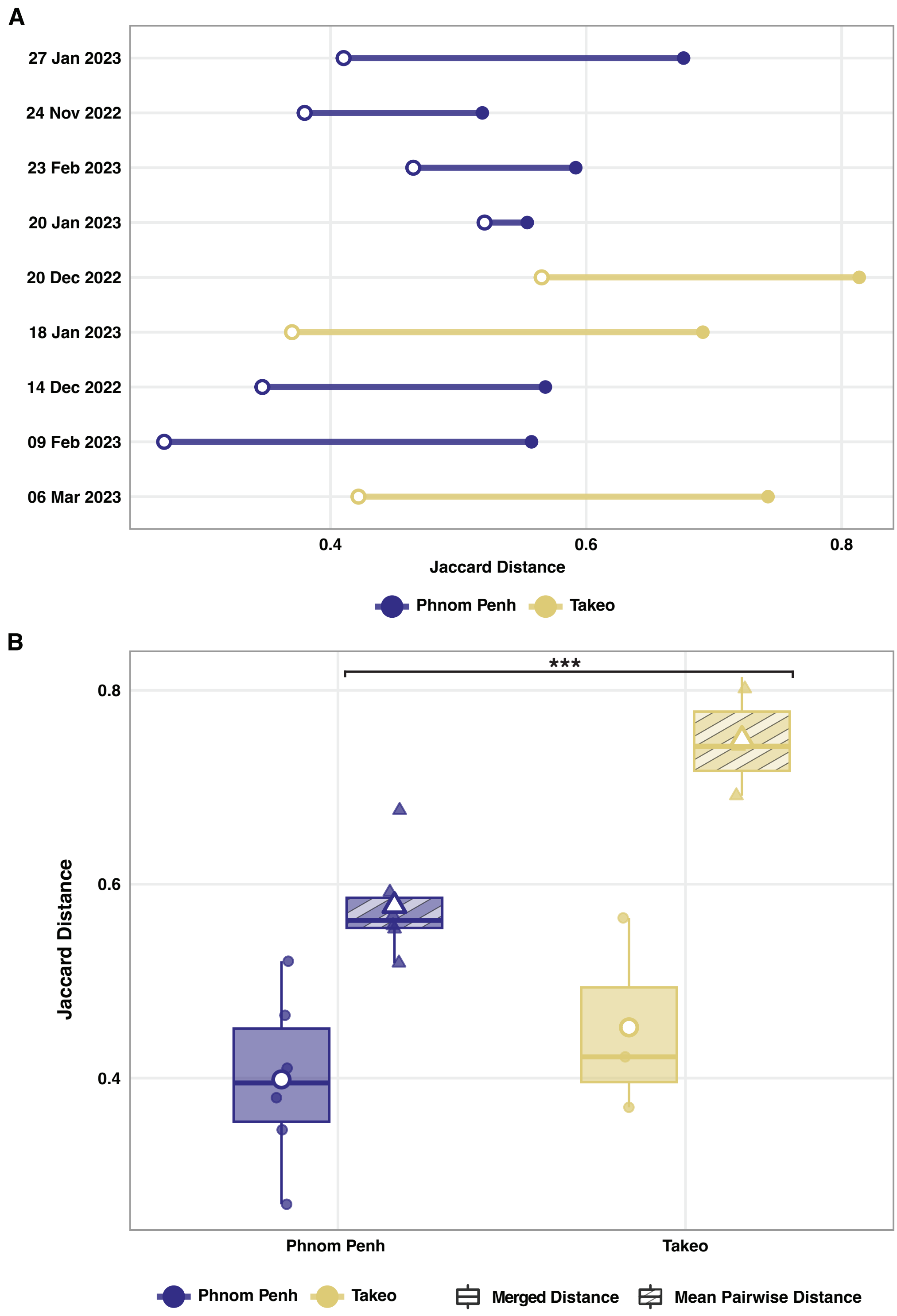


**Supplementary Figure 14: Distance between paired samples is significantly greater in Takeo than Phnom Penh.** (1), For each location and sampling date we summarised the Jaccard distance between pig and environmental samples, by merging the virus detection profile of each individual sample together. Here, a virus only had to be present in one sample for it to be reflected in the overall profile. Thus using one overall representative virus detection summary for both pig and environment at each time point we calculated the Jaccard distance between two points at each date and location. (2), we calculated the mean of each pairwise Jaccard distance between pig and individual ES types at each corresponding time point. A) At each date and location the difference between (1) merged β-diversity distances and (2) mean of all pairwise distances is shown. B) The difference between (1) and (2) is shown across all timepoints. Phnom Penh, n = 6 visits and Takeo, n = 3 visits.
