## Supplemental Figure 15 for "Air and environmental sampling expand metagenomic viral surveillance in Cambodian pig slaughterhouses"

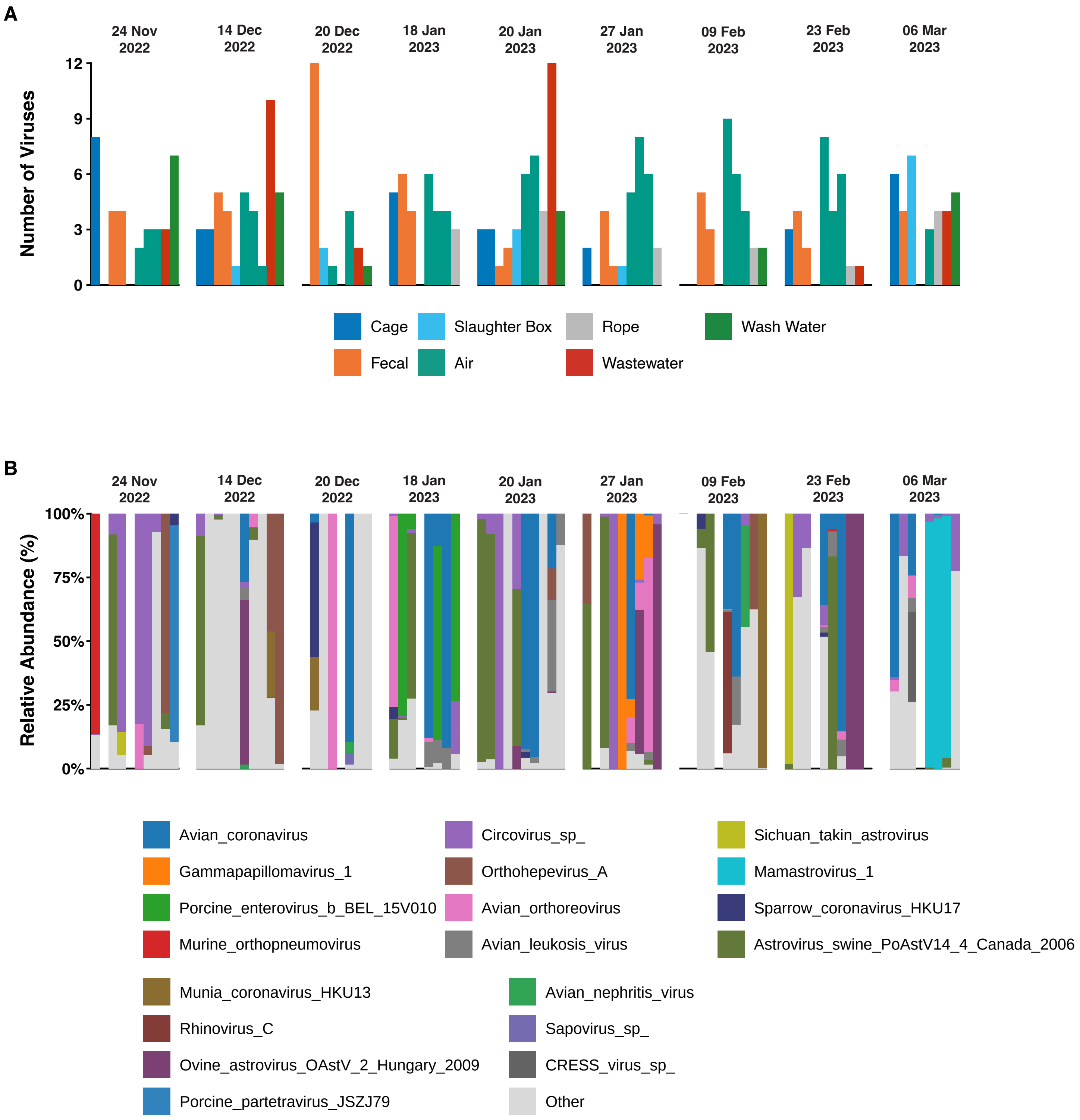


**Supplementary Figure 15: Environmental sampling can detect a broad range of non-swine viruses.** We assessed the diversity and abundance of viruses detected in the environment that were never identified through pig snout swabs at any point across the study. A) Total number of additional virus species detected in the environment broken down by sample and date. B) Stacked bar plots indicate the relative abundance (%) of the top 20 species for each sample across all sampling dates in the study.
